# Multimodal profiling and intersectional genetics reveal CA1 border inhibitory neuron diversity and postnatal emergence of retrosplenial connectivity

**DOI:** 10.64898/2026.10.29.762784

**Authors:** Natalia Martínez Vizcaíno, Nicola Thrupp, Blanca Lorente-Echeverría, Danie Daaboul, Maryam Alabi, Clara Milián Alastruey, Daniele Linaro, Yannick Fourne, Joris Vandenbempt, David Lukacsovich, Csaba Földy, Jochen Lamote, Marta Wojno, Suresh Poovathingal, Mark Fiers, Keimpe Wierda, Lynette Lim, Sara Calafate, Joris de Wit

## Abstract

Inhibitory neurons (INs) at the stratum radiatum/stratum lacunosum-moleculare border of hippocampal CA1 comprise diverse populations, including long-range-projecting neurons that innervate the retrosplenial cortex (RSCtx). However, their molecular organization and developmental connectivity remain poorly understood. Here, we combine multimodal transcriptomic profiling, intersectional genetics, and retrograde tracing to characterize CA1 border INs. We identified two major caudal ganglionic eminence-derived, *Cxcl14*-enriched populations distinguished by *Syt6/Cck*-related and *Ntng1*-expressing transcriptional profiles. We then generated a *Cxcl14*-Cre mouse line that, when combined with an IN-specific viral enhancer, provides genetic access to CA1 border INs. Retrograde labeling established that CA1 INs projecting to the RSCtx belong to the *Ntng1*-expressing population, and that this long-range projection emerges during postnatal mouse development, between postnatal days 3 and 5. Finally, analysis of genes encoding cell-surface proteins, key regulators of connectivity, revealed distinct expression profiles in RSCtx-projecting and SYT6-positive INs. Together, these findings establish a molecular and developmental framework for CA1 border IN diversity and provide a genetic strategy for investigating the assembly and function of these inhibitory circuits.

## INTRODUCTION

Connecting different neuronal cell types into functional circuits during development is critical for proper information processing in the brain [1–3]. As the principal output neurons of the hippocampal network, CA1 pyramidal neurons integrate a wide range of synaptic inputs and project to the subiculum and multiple cortical targets [4]. CA1 pyramidal neurons receive excitatory input from hippocampal CA3 region on their proximal dendrites in stratum radiatum (SR), and on their distal dendrites in stratum lacunosum-moleculare (SLM) from the entorhinal cortex. Additionally, CA1 pyramidal neurons are innervated by a diverse local array of inhibitory neurons (INs) that shape their activity and output [5–7]. Despite their central role in hippocampal information processing, the diversity and connectivity of INs in the CA1 region remain incompletely understood.

Among CA1 INs, neurons located at the SR/SLM border (hereafter referred to as CA1 border INs) remain among the least understood hippocampal IN populations. Their location places CA1 border INs at the interface between Schaffer collateral input from CA3 and perforant path input from the entorhinal cortex, positioning them as potential key players of the integration of intrahippocampal and cortical information by CA1 pyramidal neurons [4]. Despite this strategic position, CA1 border INs remain poorly characterized because of their heterogeneous nature and the limited genetic tools available for selectively accessing these neurons.

IN identities are commonly defined by a combination of molecular marker expression, electrophysiological properties, morphology, and connectivity patterns [8]. Previous studies have shown that CA1 border INs comprise a heterogeneous group of neurons, including subtypes derived from the caudal ganglionic eminence (CGE) that project their axons to either SR or SLM [9–12]. In addition, a recently described subtype extends a long-range axonal collateral to layer 1 (L1) of the retrosplenial cortex (RSCtx), where it inhibits the apical dendrites of cortical pyramidal neurons [13,14]. Single-cell sequencing analyses have identified multiple putative cell types within the CA1 border IN population [15]. However, the molecular identities of these cell types have not been definitively linked to their connectivity patterns. Furthermore, it remains unclear when subtype-specific connectivity patterns are established during development.

Here, we use Patch-seq and spatial transcriptomics to characterize the heterogeneity of CA1 border INs across developmental and adult stages. Using this transcriptomic information, we develop a *Cxcl14*-Cre driver line and establish an intersectional genetic targeting strategy for CA1 border INs. We then combine single-cell RNA sequencing of genetically labelled CA1 border INs with publicly available datasets to define the molecular identities of multiple subtypes. We find that differentially expressed genes encoding cell-surface proteins (CSPs), important regulators of axonal targeting and synaptic recognition [1–3,16], are specific molecular identifiers of CA1 border IN cell types and markers for locally projecting and long-range projecting subtypes. Finally, we focus on the long-range projecting CA1 border IN subtype, validate subtype-specific CSP markers, and determine the developmental time window during which its long-range axons reach their cortical target. Together, our findings provide a molecular and developmental framework for studying the assembly and function of CA1 border inhibitory circuits and establish the *Cxcl14*-Cre line as a valuable genetic tool for selectively accessing these neurons.

## RESULTS

### Molecular and electrophysiological heterogeneity of CA1 border INs

To characterize the heterogeneity of CA1 border INs, we first generated electrophysiological and transcriptional profiles of these neurons using Patch-seq. We performed whole-cell patch-clamp recordings from cell bodies located within +/- 100 µm of the CA1 SR/SLM border in acute sagittal brain slices containing dorsal hippocampus from young adult wild-type mice (Fig. 1A). We recorded 200 cells and characterized basic electrophysiological properties, including passive membrane properties such as resting membrane potential, as well as active properties such as rheobase current and action potential firing. Following recordings, cytosolic contents were aspirated for subsequent RNA-sequencing (RNA-seq), as previously described [17–19] (Fig. 1A). A total of 108 cells, hereby defined as Patch-seq cells, passed bioinformatic quality control (see Methods) for both transcriptional and electrophysiological analyses.

**Fig. 1.**
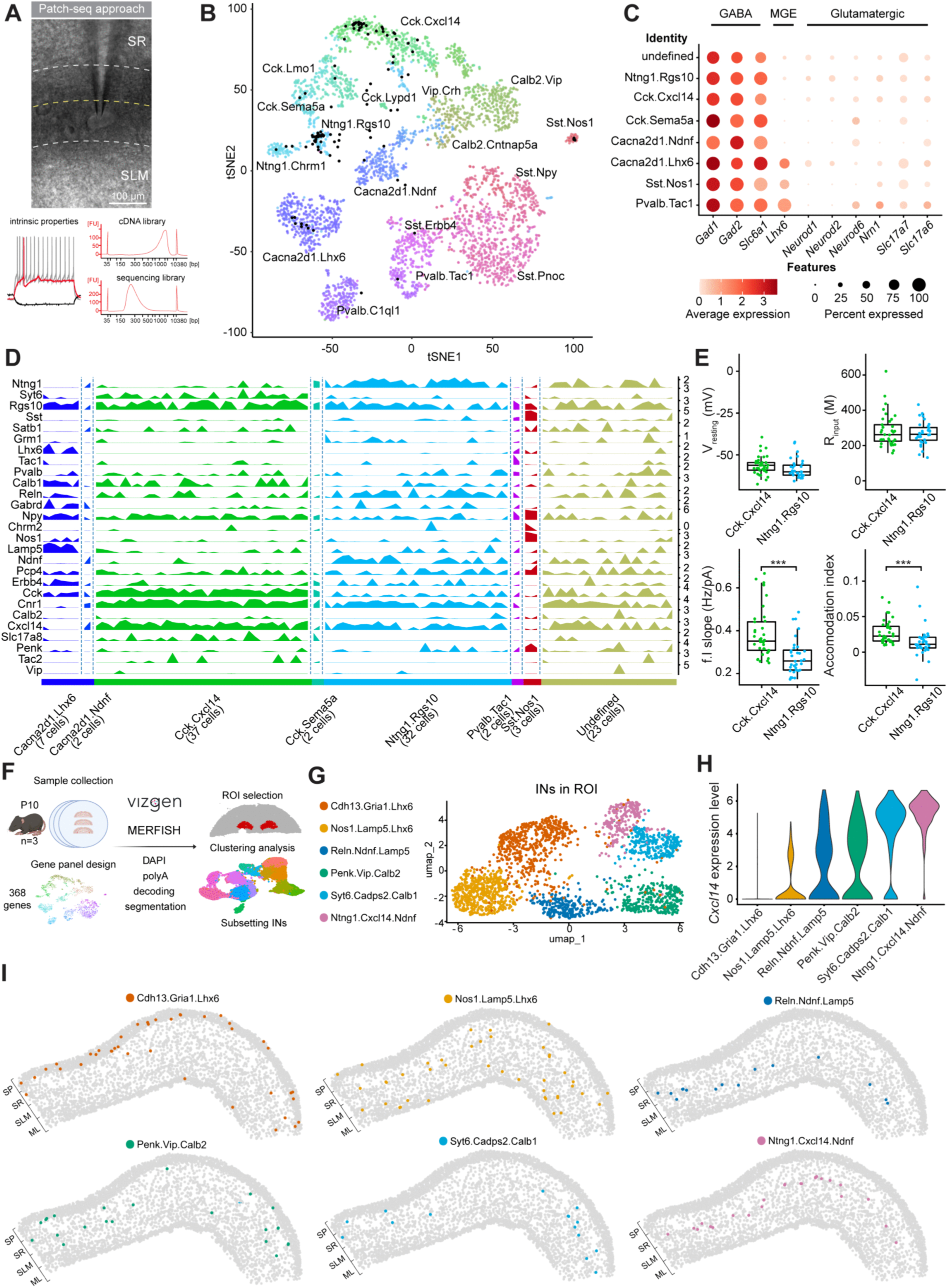
CA1 border INs express *Cxcl14* and segregate into *Ntng1*- and *Syt6*-expressing populations. (A) Schematic representation of the Patch-seq workflow used to profile intrinsic electrophysiological properties and transcriptomes of CA1 border INs. (B) Mapping of Patch-seq cells onto the Harris dataset, a reference dataset for scRNA-seq CA1 INs. Cluster labels are imported from the original dataset. (C) Dot plot showing expression of inhibitory, MGE-derived, and glutamatergic marker genes in Patch-seq cells. Dot size indicates the percentage of cells expressing each gene, and colour indicates average expression. (D) Track plot showing expression of selected marker genes across Patch-seq cells mapped onto the Harris dataset. (E) Whole-cell current clamp recordings of intrinsic properties of CA1 border INs in acute hippocampal slices from young adult wild-type mice. Box plots show resting membrane potential (V_resting_), input resistance (R_input_), frequency-current relationship (f.I slope), and accommodation index of Patch-seq cells assigned to the Ntng1.Rgs10 and Cck.Cxcl14 clusters. Boxes represent the interquartile range (IQR), horizontal lines indicate the median, and whiskers extend to the most extreme data points within 1.5 × IQR. Two-tailed unpaired t-test or Mann–Whitney test was used, depending on normality. Individual data points (***p < 0.001); Ntng1.Rgs10: n=32 cells; Cck.Cxcl14: n=37 cells. (F) Schematic representation of the MERFISH spatial transcriptomics workflow, including gene panel design, region-of-interest selection, segmentation, and IN subsetting. (G) UMAP visualization of INs from the spatial transcriptomics dataset, colored by cluster identity. (H) Violin plot showing *Cxcl14* expression across IN clusters. (I) Spatial distribution of IN clusters in the dorsal hippocampal CA1 region. Right side of image corresponds to proximal CA1. Dots represent individual cells. Abbreviations: ROI, region of interest; SP, stratum pyramidale; SR, stratum radiatum; SLM, stratum lacunosum moleculare; ML, molecular layer.

We next mapped the transcriptional profiles of the Patch-seq cells to a reference single-cell RNA-seq dataset of CA1 INs [15] (hereafter referred to as the ‘Harris dataset’), using an “out-of-sample mapping” bioinformatics pipeline [20] (Fig. 1B). Patch-seq cells mapped to seven clusters within the Harris dataset (Fig. 1B). Transcriptional analysis confirmed robust expression of GABAergic markers (*Gad1*, *Gad2*, and *Slc6a1*) and absence of glutamatergic markers (*Neurod1*, *Neurod2*, *Neurod6*, *Nrn1*, *Slc17a6*, and *Slc17a7*) across all mapped clusters (Fig. 1C).

Most Patch-seq cells mapped to CGE-derived clusters (*Lhx6*-negative): Cck.Cxcl14 (37 cells), Ntng1.Rgs10 (32 cells), Cacna2d1.Ndnf (2 cells), and Cck.Sema5a (2 cells) (Fig. 1D). A smaller number of cells mapped to medial ganglionic eminence (MGE)-derived clusters (*Lhx6*-positive), including Cacna2d1.Lhx6 (7 cells), Pvalb.Tac1 (2 cells), and Sst.Nos1 (3 cells) [21,22]. Among these MGE-derived clusters, Cacna2d1.Lhx6 cells are expected to localize near the SR/SLM border, whereas Pvalb.Tac1 and Sst.Nos1 are typically localized to stratum pyramidale (SP) and stratum oriens (SO), respectively [5,15]. An additional 23 cells could not be confidently assigned to a cluster and were classified as “undefined” (Fig. 1D).

We next analyzed the electrophysiological properties of the mapped Patch-seq clusters, focusing on the main CGE-derived populations. Cck.Cxcl14 and Ntng1.Rgs10 cells did not differ in passive membrane properties such as resting membrane potential, but Cck.Cxcl14 cells exhibited a steeper frequency-current (f/I) relationship (higher excitability) and a higher action potential accommodation index (increased spike-frequency irregularity) compared to Ntng1.Rgs10 cells (Fig. 1E and Fig. S1). Together, these results indicate that CA1 border INs are predominantly CGE-derived and comprise two principal molecular and electrophysiological populations: Cck.Cxcl14 and Ntng1.Rgs10.

### Spatial transcriptomics identifies *Syt6*- and *Ntng1*-positive CA1 border IN populations

To further characterize the spatial distribution of CA1 border INs, we performed targeted spatial transcriptomics using a custom MERFISH/Vizgen 368 gene panel based on the Harris dataset (Fig. 1F). The panel included broad cell-type markers, IN markers, and genes differentially expressed between the Cck.Cxcl14 and Ntng1.Rgs10 clusters (Table S1). In addition, genes encoding CSPs and regulators of IN type-specific connectivity patterns [23] were included in the panel. We analyzed tissue from three postnatal day 10 (P10) mice, using three sections per brain, and selected a region of interest (ROI) spanning from the CA1 SP to the dentate gyrus granule cell layer for downstream analysis (Fig. 1G).

Initial clustering of all cells within the ROI identified the expected major cell classes (Fig. S2A, B). Re-clustering of INs resolved six clusters (Fig. 1G, H), including two *Lhx6*-positive/MGE-derived populations (Cdh13.Gria1.Lhx6 and Nos1.Lamp5.Lhx6) and four *Lhx6*-negative/CGE-derived populations (Reln.Ndnf.Lamp5, Penk.Vip.Calb2, Syt6.Cadps2.Calb1, and Ntng1.Cxcl14.Ndnf) (Fig. 1H). Some clusters were broadly distributed across CA1 laminae, whereas others exhibited more spatially restricted localization patterns (Fig. 1I). The Syt6.Cadps2.Calb1 and Ntng1.Cxcl14.Ndnf clusters localized almost exclusively to the CA1 SR/SLM border (Fig. 1I and Fig. S2C). We did not observe any preference in localization along either the medial-lateral axis or rostro-caudal axes.

We next compared the expression of marker genes from the two major Patch-seq populations with the clusters from the spatial transcriptomics dataset. The SR/SLM border-localized Syt6.Cadps2.Calb1 and Ntng1.Cxcl14.Ndnf populations shared transcriptional signatures with the Cck.Cxcl14 and Ntng1.Rgs10 Patch-seq populations, respectively (Fig. S2D). Both populations robustly expressed Cxcl14 (Fig. 1D, H and Fig. S2D). Thus, the spatial transcriptomics and Patch-seq analyses consistently identified two major CA1 border IN subpopulations: a *Syt6*-positive population corresponding to the Cck.Cxcl14 subtype and a *Ntng1*-positive population. These two populations are thought to differ in connectivity, with Ntng1.Rgs10 representing a putative long-range subtype projecting to the RSCtx, and Cck.Cxcl14 comprising a heterogeneous locally projecting population [15].

We then analyzed CSP expression in our spatial transcriptomics dataset. Clustering based solely on CSP expression was sufficient to identify the same IN populations (Fig. S2E). The Syt6.Cadps2.Calb1 and Ntng1.Cxcl14.Ndnf clusters identified this way displayed the same expression pattern for Cck.Cxcl14 and Ntng1.Rgs10 marker genes (Fig. S2F), indicating that differential CSP expression is a defining feature of CA1 IN heterogeneity.

Taken together, the Patch-seq and spatial transcriptomics analyses demonstrate that CA1 border INs are heterogeneous and can be separated into at least two major molecularly defined populations at the SR/SLM border: a *Syt6*-positive/Cck-related population and a *Ntng1*-positive population.

### An intersectional genetic strategy to access CGE-derived CA1 border INs

Establishing selective genetic access to CA1 border INs has remained challenging. As shown by mRNA detection using single-molecule fluorescent in situ hybridization (smFISH, RNAscope), *Gad2* and *Cck* transcripts label broad populations of CA1 INs that include border INs, with Cck transcripts also detected in pyramidal cells.(Fig. 2A). Consequently, available Gad2-or *Cck*-Cre driver lines do not provide selective genetic access to CA1 border INs. Our Patch-seq and spatial transcriptomic analyses identified *Cxcl14* as a potential marker for CGE-derived CA1 border INs (Fig. 1I and S2C). Most Patch-seq cells mapped onto *Cxcl14*-expressing clusters in the Harris dataset (Fig. 2B; compare to Fig. 1B) [15]. smFISH for *Cxcl14* transcripts confirmed selective labeling of cells at the SR/SLM border in CA1 (Fig. 2A). We therefore generated a *Cxcl14*-Cre mouse line to establish genetic access to CA1 border INs. Using CRISPR/Cas9-mediated genome editing, we replaced the endogenous *Cxcl14* stop codon with a T2A-Cre cassette (Fig. S3 and Methods).

**Fig. 2.**
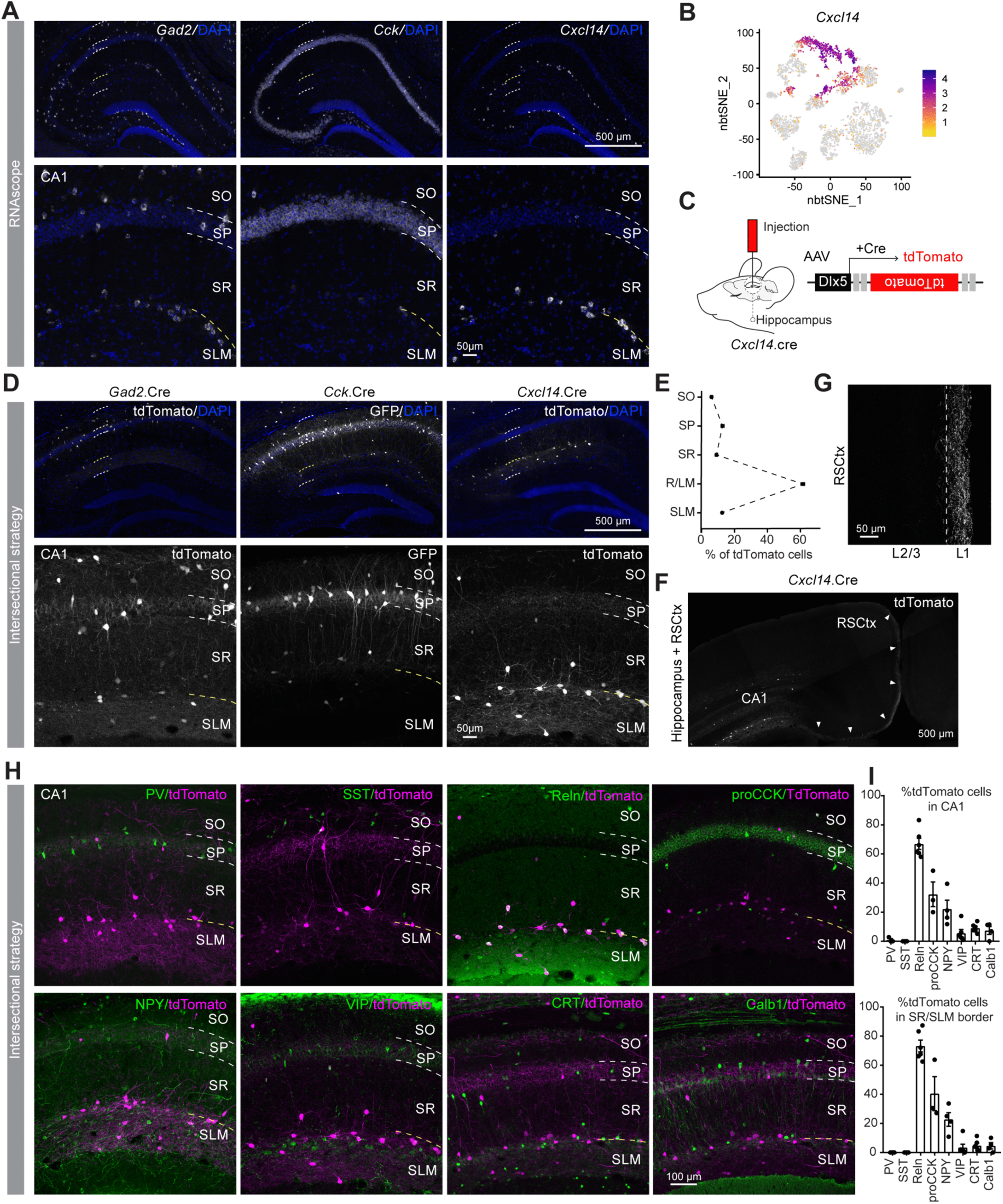
Genetic access to CA1 border INs using an intersectional *Cxcl14*-Cre and IN-specific viral enhancer strategy. (A) Coronal sections of adult mouse dorsal hippocampus (upper panel) and CA1 region (lower panel) showing expression of *Gad2*, *Cck*, and *Cxcl14* (grey). White dashed lines indicate laminar borders and yellow dashed line highlights the SR/SLM border. (B) Expression of *Cxcl14* in the Harris dataset. (C) Schematic representation of the intersectional strategy combining *Cxcl14*-Cre mice with Cre-dependent AAV under the control of the IN-specific Dlx enhancer (AAV-Dlx-FLEX-tdTomato). (D) Coronal sections of dorsal hippocampus (upper panel) and CA1 region (lower panel) showing reporter expression (grey) after injection of AAV-Dlx-FLEX-tdTomato in *Gad2*-Cre and *Cxcl14*-Cre mice, and or AAV-Dlx-FLEX-GFP in *Cck*-Cre mice. (E) Quantification of the percentage of tdTomato-positive cells (grey) across CA1 laminae following injection of AAV-Dlx-FLEX-tdTomato in *Cxcl14*-Cre mice. Graph shows mean ± SEM; n=9 animals. (F) Representative image showing tdTomato-positive axons (grey) extending from CA1 toward the RSCtx following injection of AAV-Dlx-FLEX-tdTomato in *Cxcl14*-Cre mice. (G) Representative image of tdTomato-positive axons (grey) in L1 RSCtx, signalled by arrowheads. (H-I) Coronal sections showing tdTomato reporter expression (magenta) and IHC for canonical IN markers (green) following injection of AAV-Dlx-FLEX-tdTomato in *Cxcl14*-Cre mice. (H) Representative images of co-localization and (I) quantification of co-localization in CA1 region (upper graph) and at the SR/SLM border (lower graph). Bars show mean ± SEM. Dots represent individual animals. Abbreviations: SO, stratum oriens; SP; stratum pyramidale; SR, stratum radiatum; SLM, stratum lacunosum moleculare; R/LM, border of stratum radiatum and lacunosum moleculare; PV, parvalbumin; SST, somatostatin; Reln, Reelin; proCCK, pro-cholecystokinin; NPY, neuropeptide Y; VIP, vasoactive intestinal polypeptide; CRT, calretinin; Calb1, calbindin 1.

To characterize the *Cxcl14*-Cre line, we initially crossed *Cxcl14*-Cre mice with the Cre-dependent Ai9 Rosa-CAG-LSL-tdTomato reporter line [24]. This approach resulted in abundant labeling of astrocytes in adult CA1 (data not shown), likely reflecting low but detectable *Cxcl14* expression in astrocytes (http://mousebrain.org). We therefore developed an intersectional strategy, in which an adeno-associated viral vector (AAV) expressing Cre-dependent tdTomato under the control of the inhibitory Dlx5 (mDlx5) enhancer (AAV-mDlx5-FLEX-tdTomato) was injected into the dorsal CA1 region of adult *Cxcl14*-Cre mice (Fig. 2C). This approach resulted in restricted labeling of cells in CA1 (Fig. 2D), with 62% of tdTomato-positive neurons localized to the CA1 SR/SLM border (Fig. 2E). Importantly, the same intersectional approach with *Gad2*-or *Cck*-Cre lines labeled neurons in CA1 more broadly (Fig. 2D).

In addition to labeling neuronal somata in CA1 in *Cxcl14*-Cre mice, the intersectional approach revealed a dense tdTomato-positive axon bundle projecting to L1 of the RSCtx (Fig. 2F, G). This observation confirms that a subpopulation of CA1 border INs labeled by this strategy extends long-range projections to the RSCtx, consistent with a previous report [14].

We next characterized the identity of tdTomato-labeled cells in *Cxcl14*-Cre mice injected with AAV-mDlx5-FLEX-tdTomato using immunohistochemistry (IHC) for canonical IN markers. Approximately 70% of tdTomato-positive neurons at the CA1 border were positive for Reelin, whereas 40% expressed proCCK and 20% expressed neuropeptide Y (NPY) (Fig. 2H, I). A smaller fraction (<10%) of CA1 border INs expressed Calbindin, Calretinin, or VIP. In contrast, Parvalbumin and somatostatin were absent from tdTomato-positive border INs (Fig. 2H, I). Together, these findings demonstrate that the intersectional strategy combining the *Cxcl14*-Cre line with an inhibitory enhancer-driven Cre-dependent reporter provides genetic access to CA1 border INs and preferentially labels Reelin-positive, CGE-derived neurons within this population.

### scRNA-seq characterization of genetically-labeled postnatal CA1 border INs

To further characterize CA1 border INs labeled using the intersectional strategy in *Cxcl14*-Cre mice, we performed single-cell RNA-sequencing (scRNA-seq) analysis. We selected P3, P5, and P10 as analysis timepoints spanning key stages of postnatal IN development, including continued neuronal migration and laminar positioning, early synaptogenesis, and developmental programmed cell death [9,22,25]. To selectively label CA1 border INs during this developmental window, we performed intracerebroventricular (ICV) injections of AAV-hDlx-FLEX-tdTomato in *Cxcl14*-Cre pups at P0. This strategy resulted in robust tdTomato expression in CA1 border INs at P3, P5, and P10 (Fig. 3A).

**Fig. 3.**
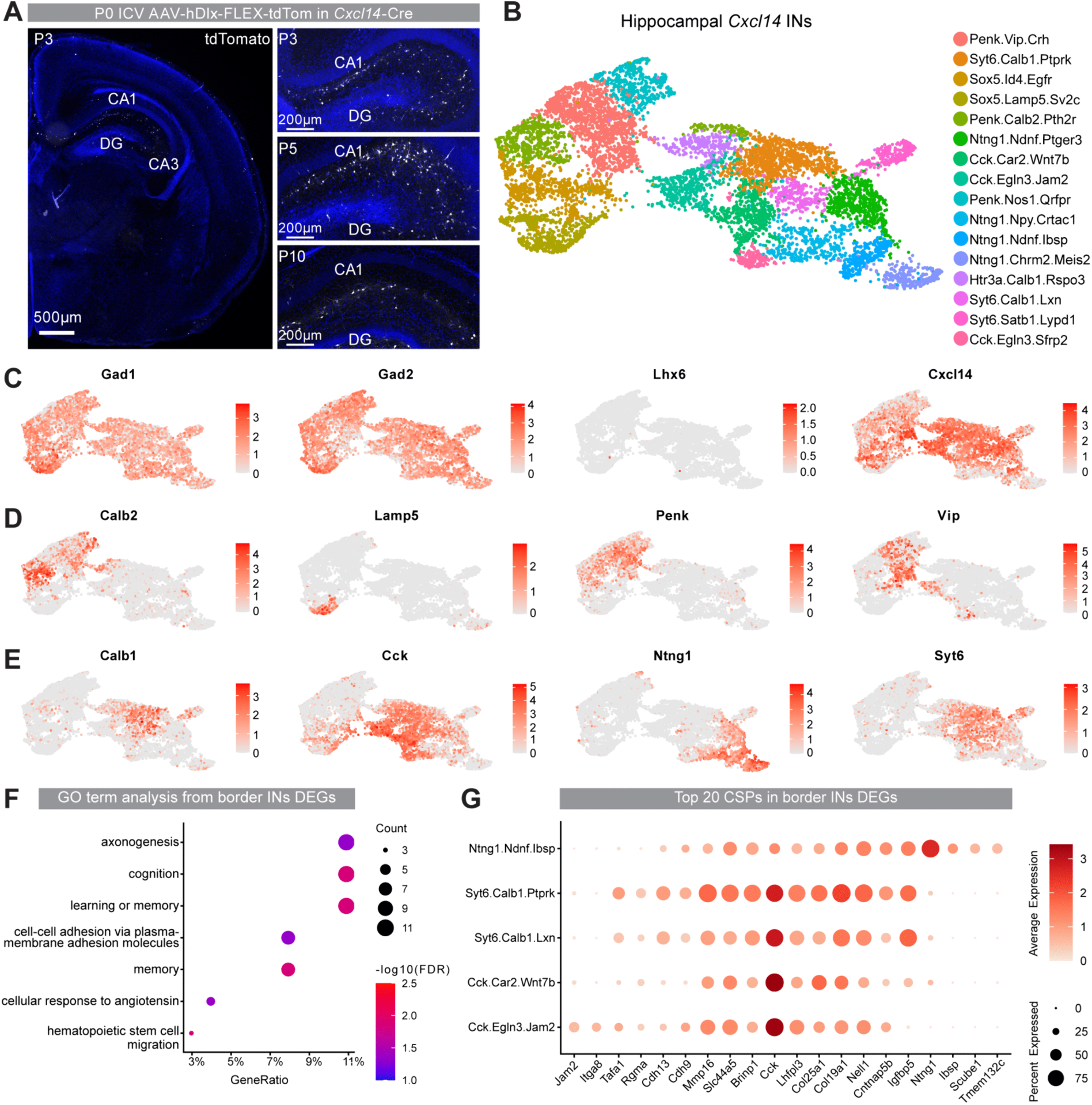
scRNA-seq characterization of postnatal *Cxcl14*-expressing hippocampal INs. (A) Representative images from P3, P5, and P10 dorsal hippocampus following P0 intracerebroventricular (ICV) injection of AAV-Dlx-FLEX-tdTomato in *Cxcl14*-Cre mice. (B) UMAP visualization of the postnatal *Cxcl14*-lineage hippocampal IN scRNA-seq dataset after quality control, coloured by cluster identity. (C) Feature plots showing expression of IN and *Cxcl14*-lineage markers, colour indicates average expression. (D) Feature plots showing markers enriched in clusters located on the left side of the UMAP, colour indicates average expression. (E) Feature plots showing markers enriched in clusters located on the right side of the UMAP, colour indicates average expression. (F) GO enrichment analysis of DEGs across CA1 border IN clusters. Dot size indicates the number of genes associated with each GO term, and colour indicates enrichment significance, shown as −log10 false discovery rate (FDR)-adjusted p-value. (G) Dot plot showing top 20 CSP genes, based on lowest p-value, in clusters corresponding to CA1 border INs based on mapping to the Harris dataset. Dot size indicates the percentage of cells expressing each gene, and colour indicates average expression.

Following tissue dissociation, tdTomato-positive cells were isolated by fluorescence-activated cell sorting (FACS) and processed using the 10X genomics platform (see Methods) [26]. To maximize cell recovery, whole hippocampi were collected for analysis; accordingly, we refer to this dataset as “*Cxcl14* hippocampal INs” to indicate that labeled and sorted cells are not confined to the CA1 SR/SLM border. After removal of low-quality cells and contaminating cell types (see Methods), the final dataset consisted of 8.187 cells, including 1.244 cells from P3, 5.584 cells from P5 cells, and 1.359 cells from P10 samples.

Unsupervised clustering analysis based on highly variable genes identified 16 transcriptionally distinct populations (Fig. 3B). Batch correction using Harmony [27] effectively minimized sample-specific structure, as all clusters contained cells from each developmental stage and UMAP visualization showed broad intermixing across samples (Fig. S4A, B). In addition, the number of detected genes and percentage of mitochondrial genes were broadly comparable across clusters (Fig. S4C, D), indicating that clustering was not driven by cell quality.

Cluster identities were assigned based on known markers and differentially expressed genes (DEGs). UMAP visualization revealed broad expression of the inhibitory markers *Gad1* and *Gad2* across clusters, confirming the GABAergic identity of the isolated cells (Fig. 3C). *Cxcl14* was expressed throughout the dataset, whereas expression of the MGE-associated marker *Lhx6* was minimal, consistent with preferential labeling of CGE-derived populations by the intersectional strategy (Fig. 3C). Reference mapping to the Allen Brain taxonomy confirmed that the dataset was dominated by CGE-derived inhibitory subclasses, including Vip-, Sncg-, Lamp5-, and Retrohippocampal region-Cortical amygdalar area (RHP-COA)/Ndnf-related populations (Fig. S5A, B) [28].

To determine which developmental populations corresponded to the adult Cck.Cxcl14 and Ntng1.Rgs10 populations identified in the Patch-seq analysis, we next performed label transfer using the Harris dataset (Fig. S5C, D) [15]. Several developmental clusters mapped to the adult Cck.Cxcl14 population, including Cck.Car2.Wnt7b, Cck.Egln3.Jam2, Syt6.Calb1.Ptprk and Syt6.Calb1.Lxn (Fig. S5C, D), clusters that are located adjacent to each other in our UMAP (Fig. 3B). This finding is consistent with the heterogeneity previously described for adult Cck.Cxcl14 cells by Harris et al. (Table S2) and suggests that this population comprises transcriptionally distinct subtypes. This heterogeneity is further illustrated by the fact that most of these developmental clusters expressed *Syt6*, with the exception of the Cck.Egln3.Jam2 cluster (Fig. 3E), consistent with the molecular diversity encompassed within the adult Cck.Cxcl14 class.

In contrast, label transfer identified a more discrete *Ntng1*-enriched transcriptional component (Fig. S5C, D). Among these clusters, Ntng1.Ndnf.Ibsp showed unique correspondence to adult Ntng1.Rgs10, the putative RSCtx-projecting CA1 border IN subtype as defined by Harris et al. [15]. Additional *Ntng1*-expressing clusters mapped to related but distinct adult populations. For example, Ntng1.Chrm2.Meis2 was readily identified as a trilaminar-cell-related population based on its selective Chrm2 expression and was therefore unlikely to correspond to CA1 border INs [29,30]. Together, these results indicate that *Ntng1* expression marks a broader set of hippocampal IN populations, only a subset of which closely corresponds to the putative long-range-projecting Ntng1.Rgs10 population.

### scRNA-seq reveals differentially expressed CSPs among developing CA1 border INs

Having associated Ntng1.Ndnf.Ibsp, Cck.Car2.Wnt7b, Cck.Egln3.Jam2, Syt6.Calb1.Ptprk, and Syt6.Calb1.Lxn with CA1 border INs based on Harris label transfer and Patch-seq enrichment, we next examined their DEGs in more detail. First, to assess the functional significance of these transcriptional differences, we performed Gene Ontology enrichment analysis. Enriched categories included axonogenesis and cell-cell adhesion via plasma-membrane adhesion molecules, as well as broader terms such as cognition and learning or memory (Fig. 3F). Together, these findings suggest that molecular differences among hippocampal *Cxcl14*-positive populations are associated with gene programs linked to axon targeting and synaptic partner selection.

We then focused on genes encoding CSPs as regulators of connectivity, which revealed several distinct patterns. Some CSPs were shared across multiple CA1 border IN clusters, including *Col19a1*, *Col25a1*, and *Igfbp5*. However, we also identified CSPs that were selective for specific clusters, including *Ibsp*, *Scube1*, and *Tmem132c* in Ntng1.Ndnf.Ibsp, and *Jam2* and *Itga8* in Cck.Egln3.Jam2 (Fig. 3G). These findings suggest that CSP expression is organized at multiple levels, with some molecules marking broader subclass relationships and others distinguishing individual transcriptomic populations that may differ in connectivity patterns.

Lastly, we examined the temporal expression patterns of these CSP genes (Fig. S6A-C). A previous study suggested that postnatal time points before P10 are particularly relevant for studying CSP expression dynamics and their contribution to connectivity, as the expression of some CSPs becomes undetectable after P10 [23]. *Ibsp* appeared to follow this pattern, showing a tendency toward lower expression at later postnatal ages in the Ntng1.Ndnf.Ibsp cluster (Fig. S6A). Other CSP genes, such as *Cdh13*, followed a trend towards the opposite pattern, with higher expression at later ages (Fig. S6B). Although several CSP genes were expressed across multiple CA1 border IN clusters, their temporal expression patterns often differed between clusters. For example, *Cntnap5b* expression showed a tendency to increase with postnatal age in Cck.Egln3.Jam2 cells, while showing the opposite tendency in the other CA1 border IN clusters (Fig. S6C). Together, these observations suggest that CSPs in CA1 border INs are expressed in subtype-specific and temporally defined patterns.

### RSCtx-projecting CA1 border INs express *Ntng1* and are SYT6-negative

To begin linking molecular identity of CA1 border IN populations to specific connectivity patterns, we focused on the two main subpopulations identified by Patch-seq, spatial transcriptomics, and scRNA-seq, marked by *Ntng1* and *Syt6*, respectively. Taking advantage of their distinct connectivity pattern, we restricted this analysis to RSCtx-projecting CA1 border INs, a subtype hypothesized by Harris et al. to correspond to the adult Ntng1.Rgs10 cluster [15]; the Ntng1.Ndnf.Ibsp cluster in our developmental *Cxcl14* hippocampal IN dataset (Fig. S5D). A previous study showed that RSCtx-projecting CA1 INs at the SR/SLM border can be retrogradely labelled by injecting retrobeads into layer 1 (L1) of the RSCtx [14].

We performed retrobead injections into RSCtx L1 of adult mice and processed tissue for *Ntng1* smFISH and SYT6 immunohistochemistry (IHC) two days later (Fig. 4A). Retrobead-labeled cells at the CA1 SR/SLM border were *Ntng1*-positive but SYT6-negative (Fig. 4B), confirming that RSCtx-projecting CA1 border INs belong to the *Ntng1*-expressing subgroup. SYT6-positive neurons, on the other hand, were not retrogradely labeled from the RSCtx, consistent with their proposed identity as locally projecting INs.

**Fig. 4.**
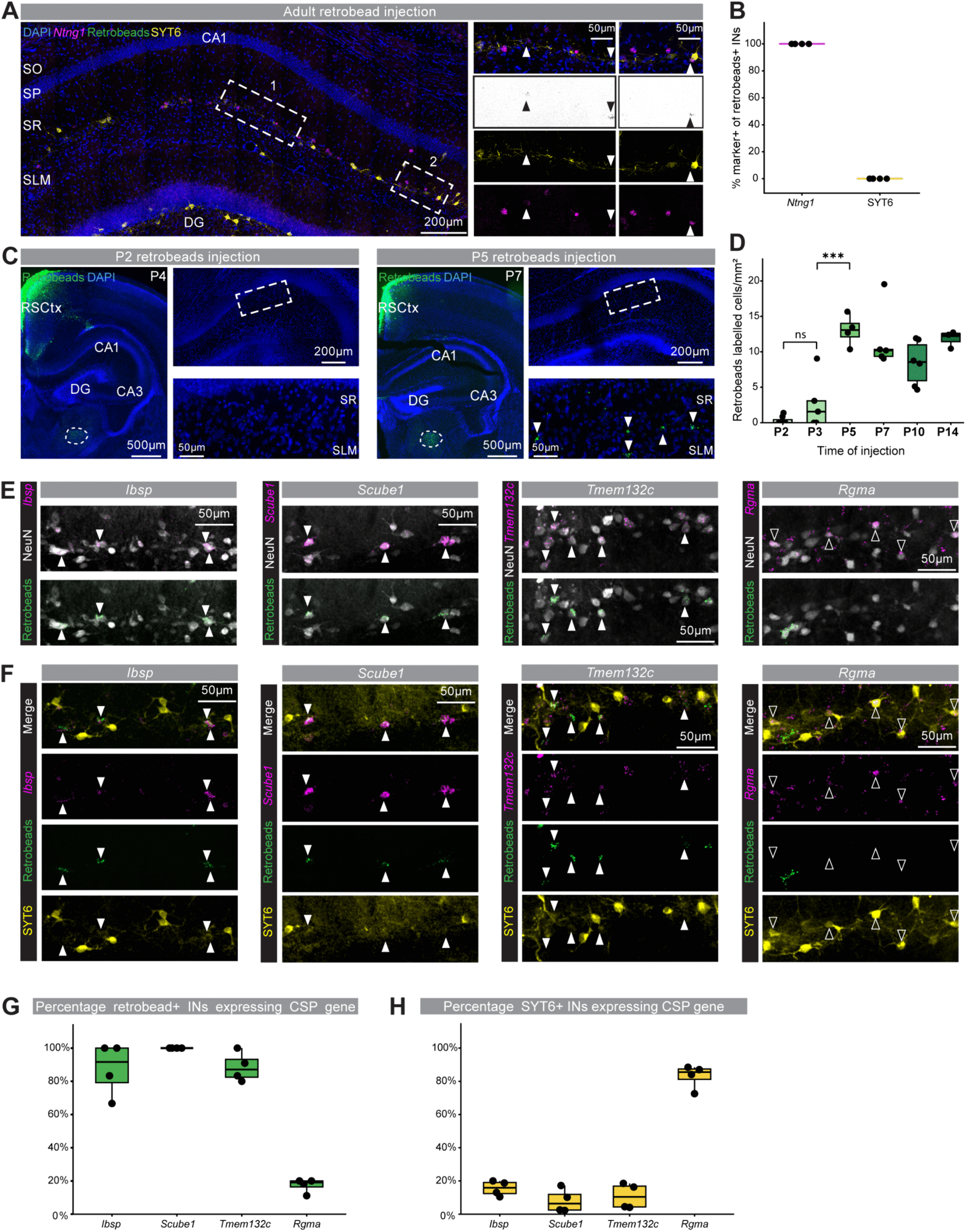
Molecular and developmental characterization of RSCtx-projecting CA1 border INs. (A) Hippocampal section (left) and CA1 SR/SLM border crops (right) showing *Ntng1* expression (magenta), SYT6 immunoreactivity (yellow), and retrobead labeling (green) two days after retrobead injection in RSCtx L1 in adult mice. (B) Quantification of *Ntng1* expression and SYT6 immunoreactivity in retrobead-labeled cells, two days after retrobead injection in RSCtx L1 in adult mice. Dots represent individual animals. (C) Representative images showing CA1 retrobead labeling (green) two days after retrobead injection in RSCtx L1 at P2 (left) or P5 (right). Dotted boxes represent from which the zoomed SR/SLM border crop was obtained. Arrowheads point to retrobead labelled cells. (D) Quantification of retrobead-labeled cells (green) normalized to CA1 SR/SLM area two days after RSCtx L1 injection at different postnatal ages. Boxes represent the interquartile range (IQR), horizontal lines indicate the median, and whiskers extend to the most extreme data points within 1.5 × IQR. Dots represent individual animals. One-way ANOVA was used to determine statistical significance, followed by Tukey’s multiple comparisons test (*** p < 0.001; n.s., not significant). (E) CSP gene expression (magenta) and retrobead labeling (green) at the CA1 SR/SLM border together with NeuN immunohistochemistry (grey), two days after P5 retrobead injection in RSCtx L1. Filled arrowheads indicate cells that display colocalization between CSP expression and retrobead labeling; empty arrowheads indicate cells that display colocalization between CSP expression and SYT6 labeling. (F) Shows the same fields of view from panel E with different combinations of channels. CSP gene expression (magenta), SYT6 immunoreactivity (yellow), and retrobead labeling (green) at the CA1 SR/SLM border two days after P5 retrobead injection in RSCtx L1. Filled arrowheads indicate cells that display colocalization between CSP expression and retrobead labeling; empty arrowheads indicate cells that display colocalization between CSP expression and SYT6 labeling. (G) Boxes represent the interquartile range (IQR), horizontal lines indicate the median, and whiskers extend to the most extreme data points within 1.5 × IQR. Quantification of the percentage of retrobead-positive INs expressing each CSP gene. Dots represent individual animals. (H) Quantification of the percentage of SYT6-positive INs expressing each CSP gene. Boxes represent the interquartile range (IQR), horizontal lines indicate the median, and whiskers extend to the most extreme data points within 1.5 × IQR. Dots represent individual animals.

### Long-range projections from CA1 border INs reach RSCtx layer 1 between P3 and P5

To determine when the long-range projection from CA1 border INs reaches the RSCtx target, we performed retrobead labeling across postnatal development (Fig. 4C). Retrobeads were injected into RSCtx L1 and animals were perfused two days later to allow retrograde transport. Following RSCtx injection at P2, virtually no retrobead-labeled neurons were detected in the CA1 SR/SLM border (Fig. 4C, D; Fig. S7). Injections at P3 yielded mixed results: some animals displayed a small number of labeled neurons, whereas others showed none (Fig. 4D). From P5 onward, RSCtx retrobead injections consistently resulted in labeling of cells at the CA1 SR/SLM border (Fig. 4C, D), suggesting that axons from long-range-projecting CA1 border INs reach RSCtx L1 between P3 and P5, establishing their target innervation by P5.

### Long-range projecting CA1 border INs express a specific combination of CSPs

To further characterize the link between molecular identity and connectivity pattern, we selected the top three differentially expressed CSP genes from the scRNA-seq Ntng1.Ndnf.Ibsp cluster: Integrin Binding Sialoprotein (*Ibsp*), Signal Peptide, CUB and EGF-like Domain Containing 1 *(Scube1*), and Transmembrane Protein 132c (*Tmem132c*). We also included a CSP gene enriched in the *Syt6* clusters, Repulsive Guidance Molecule A (*Rgma*) [31]. To confirm whether these genes were enriched in long-range projecting or *Syt6-*positive CA1 border INs, retrobeads were injected in RSCtx L1 at P5. Tissue was collected at P7 and smFISH was performed for each of the CSP gene in combination with SYT6 IHC.

The four CSP genes were expressed in CA1 border INs, as shown by gene expression colocalization with NeuN-labelled cells at the SR/SLM border (Fig.4E, Fig. S8). They showed different expression patterns in the hippocampus, with *Scube1* displaying the most border IN-specific pattern while the other CSPs were also expressed in pyramidal neurons and/or non-neuronal cell types (Fig. S8B). In agreement with the scRNA-seq data, *Ibsp*, *Scube1*, and *Tmem132c* were enriched in CA1 border INs that are labelled with retrobeads (Fig. 4F, G), while showing minimal colocalization with SYT6-positive INs (Fig. 4G). *Rgma* mRNA was enriched in SYT6-positive CA1 border INs that are not labeled with retrobeads (Fig 4F, H). Taken together, these results identify the Ntng1.Ndnf.Ibsp cluster as RSCtx-projecting CA1 border INs, reveal that their long-range projection emerges early in postnatal development, and establish that these neurons express a specific combination of CSP genes.

## DISCUSSION

In this study, we combined Patch-seq, spatial transcriptomics, intersectional genetics, and developmental scRNA-seq to characterize the molecular diversity of CA1 border INs. We identified molecularly distinct local- and long-range-projecting populations and showed that long-range projections to the RSCtx emerge during a restricted postnatal developmental window. Our data further revealed that long-range-projecting INs express a distinct combination of CSPs, suggesting that subtype-specific molecular programs contribute to the establishment of long-range inhibitory connectivity.

A central finding of this study is that INs located at the CA1 SR/SLM border do not constitute a homogeneous population. Across datasets, we consistently identified two major CGE-derived and *Cxcl14*-enriched populations: a *Syt6*.*Cck*-related population and a *Ntng1*-expressing population corresponding to putative RSCtx-projecting INs. By combining transcriptomic characterization with retrograde labeling, we linked the molecular identity of the *Ntng1*-positive population to CA1 border INs projecting to layer 1 of the RSCtx. In contrast, SYT6-positive INs were not retrogradely labelled from the RSCtx, indicating that they represent a separate population that does not share this projection target. Importantly, our transcriptomic analyses further indicate that the *Syt6*- and *Ntng1*-positive populations are themselves heterogeneous.

Our developmental scRNA-seq dataset suggests that RSCtx-projecting CA1 border INs are embedded within a broader and heterogeneous group of *Ntng1*-expressing hippocampal INs. This interpretation is consistent with the greater number of *Ntng1*-positive cells than retrobead-positive cells observed at the SR/SLM border, although incomplete retrograde labeling likely also contributes to this difference. Combining retrobead labelling with smFISH confirmed that the Ntng1.Rgs10 cluster in the Harris dataset [15] and Ntng1.Ndnf.Ibsp in our developmental *Cxcl14* hippocampal IN dataset correspond to the RSCtx-projecting CA1 border INs. Beyond this projection-defined population, our dataset identified several additional *Ntng1*-expressing populations with distinct identities. Of these, Ntng1.Ndnf.Ptger3 likely represents another border-associated population. This cluster mapped predominantly to the Ntng1.Synpr population within the Harris dataset, which was not recovered in our Patch-seq mapping, but also mapped to the 0193 RHP-COA Ndnf Gaba_1 supertype in the Allen Brain Atlas. The spatial distribution of this supertype is restricted to the CA1 SR/SLM border (Table S2), suggesting a border-associated identity. By contrast, the anatomical identity of Ntng1.Npy.Crtac1 remains less clear, because its corresponding Allen supertype is broadly distributed throughout the hippocampus, with only a few cells located at the SR/SLM border. Finally, selective Chrm2 expression identifies Ntng1.Chrm2.Meis2 as a trilaminar-cell-related population that is not located at the CA1 border [29,30]. Thus, *Ntng1* expression does not uniquely identify RSCtx-projecting CA1 border INs but instead marks several molecularly and anatomically distinct hippocampal IN populations.

The presence of *Ntng1* expression across several hippocampal IN populations also raises the possibility that it is associated more broadly with extrahippocampal projection identity. Trilaminar cells are located in SO and project both across CA1 laminae and to the subiculum [29,30]. Harris et al. similarly proposed that *Ntng1* may mark CA1 INs projecting to the subiculum [15]. Axons of RSCtx-projecting CA1 border INs similarly pass through the subiculum en route to the RSCtx. *Ntng1* expression may therefore be associated with the capacity of CA1 INs to extend axons beyond the hippocampus proper, rather than with projection to a specific target. Of note, Lrrc4c, which encodes NGL-1, the binding partner of Netrin-G1, is enriched in the subiculum [32], whereas NGL-1 protein is preferentially localized to the distal dendrites of cortical pyramidal neurons in layer 1 [33]. Although these observations remain correlative, they point to the Netrin-G1/NGL-1 interaction as a candidate molecular mechanism contributing to the organization of extrahippocampal inhibitory projections.

Our developmental scRNA-seq analysis also revealed broader *Cxcl14* expression than expected from adult datasets and from characterization of the *Cxcl14*-Cre line in adulthood. Several developmental clusters expressed *Vip* and/or *Calb2*, despite the limited colocalization of these markers with INs labelled by our intersectional strategy (Fig. 2H, I). Moreover, whereas *Cxcl14* expression was restricted to a single Vip subtype in the Harris dataset, our *Cxcl14*-expressing clusters mapped across multiple Vip populations. The Ntng1.Chrm2.Meis2 cluster provided a further example of this developmental breadth. This population corresponds to trilaminar cells, which lack detectable *Cxcl14* expression in adult scRNA-seq datasets [15,29], suggesting that *Cxcl14* expression in these cells is transient and developmentally restricted. Together, these findings indicate that *Cxcl14* marks a broader and more dynamic range of CGE-derived IN populations during early postnatal development than is apparent from adult transcriptomic datasets.

We found that clustering based exclusively on CSP gene expression was sufficient to distinguish CA1 border IN populations in our spatial transcriptomic dataset. Together with the enriched GO terms identified in our developmental scRNA-seq dataset, including “axonogenesis” and “cell-cell adhesion via plasma membrane molecules”, this finding suggests that CSP composition is a core feature of CA1 border IN identity. By contrast, a Patch-seq analysis of morphologically distinct PV-positive INs in adult CA1 found largely homogeneous CSP profiles across cells targeting the somata, axon initial segments, or dendrites of CA1 pyramidal neurons [34]. Several factors may account for this difference. First, our custom CSP gene panel was designed based on the Harris dataset and may therefore have increased the sensitivity for detecting differences among CA1 border IN populations. Second, our spatial transcriptomics analysis examined a developmental time point at which differences in CSP expression may be more pronounced. Finally, CA1 border INs may encompass greater molecular and cellular diversity than the morphological subtypes examined within the PV-positive IN class.

We identified *Ibsp*, *Scube1*, and *Tmem132c* as selectively enriched in RSCtx-projecting CA1 border INs, whereas *Rgma* characterized the SYT6-positive population. The GPI-anchored protein RGM is of particular interest because of its well-established role in restricting neurite outgrowth through its interaction with neogenin [31,35]. Its expression in the putative locally projecting SYT6-positive population is therefore consistent with a possible role in limiting axonal extension. By contrast, SCUBE1 is a secreted or membrane-associated glycoprotein implicated in vascular biology and BMP signalling [36,37]. IBSP (integrin-binding bone sialoprotein) is an extracellular matrix protein studied primarily in bone biology [38]. The expression of these molecules suggests that extracellular matrix interactions and vascular-associated signalling may contribute to the development of long-range projecting INs. TMEM132c is a large transmembrane protein proposed to link the extracellular environment to the intracellular actin cytoskeleton [39,40]. Studies in *C. elegans* have implicated related proteins in cell motility and neuronal morphology, raising the possibility that TMEM132c regulates axonal growth or structural maturation [41]. Although the functions of these molecules in CA1 border INs remain unknown, their differential expression suggests that distinct CSP combinations may contribute to the establishment of local and long-range inhibitory connectivity.

The enrichment of *Cxcl14* in CA1 border INs enabled us to generate a new *Cxcl14*-Cre mouse line that provides genetic access to *Cxcl14*-expressing INs in development and adulthood. Notably, *Cxcl14* has also been used to target CGE-derived INs in other brain regions. Iannone et al. recently used a *Cxcl14*-GFP mouse line to characterize cortical L1 INs [42]. This distribution is consistent with the labeling pattern we observe in cortical regions following neonatal ICV injections of AAV-hDlx-FLEX-tdTomato in *Cxcl14*-Cre mice. Together with our developmental scRNA-seq analysis, these findings indicate that *Cxcl14*-Cre provides access to a broad class of CGE-derived INs both within and outside the hippocampus.

Our developmental scRNA-seq data further indicate that *Cxcl14* expression is broader and more dynamic during early postnatal development than in adulthood. This transient expression raises the possibility that *Cxcl14* contributes to early developmental processes such as migration or axon extension. Consistent with this possibility, CXCL14 is a chemokine reported to interact with CXCR4 [43], a receptor with a well-established role in IN migration mediated by CXCL12 signaling [44,45]. However, deletion of *Cxcl14* in cortical layer 1 INs did not alter their laminar positioning, suggesting that the function of CXCL14 is not restricted to migration and may depend on cellular and developmental context [42]. The molecular framework established here, together with the *Cxcl14*-Cre line, provides a valuable resource for investigating how CGE-derived hippocampal INs acquire subtype-specific connectivity patterns and integrate into hippocampal and cortical circuits.

## METHODS

### Animals

Mice were maintained in a specific pathogen–free facility under standard housing conditions with continuous access to food and water. All animal experiments were conducted according to KU Leuven guidelines and approved by the KU Leuven Ethical Committee for Animal Experimentation (ECD projects: P183/2017, P175/2021). Wild-type C57BL/6J mice originated from Charles River.

#### Cxcl14-Cre mouse line

The *Cxcl14*-Cre mouse line was generated by Cyagen/Taconic. Using CRISPR/Cas9, the stop codon from the Cxcl14 gene was replaced with a T2A-Cre cassette. gRNA sequence (matching forward strand of gene) was: CCCTATTCTTCGTAGACCCTGGG.

#### Gad2-Cre mouse line

The Gad2tm2(cre)Zjh/J / Gad2-IRES-Cre line (stock 010802) was obtained from JAX.

#### Cck-Cre mouse line

The Ccktm1.1(cre)Zjh/J / CCK-ires-Cre line (stock 012706) was obtained from JAX.

### Plasmids

pAAV-mDlx5-FLEX-tdTomato was generated using the FLEX-tdTomato fragment from pAAV-CAG-FLEX-tdTomato-WPRE (Lab of Synapse Biology) and the backbone from pAAV-mDlx-GFP-Fishell-1 (Addgene #83900). pRep/Cap (AAV2/9; Addgene, 112865) and pΔF6 (Addgene, 112867) plasmids were used to produce AAVs.

### AAV generation

High-titer AAV production and purification were carried out as previously reported [46]. Briefly, six plates of 80% confluent HEK cells were transfected with 20 μg of pDelta F6 plasmid, 10 μg of pRepCap 2/9 plasmid, and 10 μg of AAV genome plasmid per plate using polyethylenimine transfection. Three days later, cells were collected by scraping and centrifugation. Lysates were treated with benzonase nuclease to remove cellular DNA. Lysates were loaded onto an iodixanol gradient (60, 40, 25, and 15%) and ultracentrifuged for 1 hour 40 mins at 50,000 rpm at 12°C. The 40% fraction containing the purified AAV was carefully collected, desalted, and concentrated on a 4-ml Amicon column (Sigma-Aldrich). AAV purity was tested by silver staining using the ProteoSilver Silver Stain Kit (Sigma-Aldrich, PROTSIL1-1KT).

### Stereotactic injections

#### CA1 region of the hippocampus

Mice were anesthetized with 5% isoflurane and placed in a mouse stereotact (KOPF) on a hot plate kept at 37°C. During the rest of the procedure, around 2% isoflurane was constantly administered, depending on animal breathing, monitored by eye. After disinfecting the mouse’s head, local anesthesia was administered by a subcutaneous injection with 100 μl of lidocaine (1% xylocaine). An incision was made in the skin to reveal the skull. One unilateral injection was performed; coordinates to target CA1 from bregma: Anterior-Posterior (AP):2.2, Medial-Lateral (ML):1.7, Dorsal-Ventral (DV):-1.7 from the surface of the skull. Two hundred nanoliters of AAV-mDlx5-FLEX-tdTomato (produced in house) was injected using a Nanoject III (Drummond) at 5 nl/s. After 5-min recovery, the capillary was pulled out slowly. The incision was stitched with surgical glue (Millpledge Veterinary). After 6 hours, their health was examined, and mice were injected with buprenorphine (0.1 mg/kg).

#### RSCtx

For targeting of RSCtx in adult, mice were anesthetized and manipulated as previously described. In this case, coordinates were as follows, from bregma: AP:1.4 and 2 ML:0.1 DV:0.3,0.6,0.9, and 50nl of Green Retrobeads™ IX (Lumafluor) were injected per DV coordinate.

For targeting of RSCtx from P3 to P14, bevelled capillaries were used. ML coordinate was determined by eye, right next to the midline blood vessels. The rest of the coordinates were taken from lambda. As described before, 50nl of retrobeads were injected per DV coordinate.

P14 coordinates: AP:1.5 and 2, DV:0.4 and 0.8

P10 coordinates: AP:1.3 and 1.8, DV:0.4 and 0.8

P7 coordinates: AP:1.1 and 1.5, DV:0.35 and 0.7

P5 coordinates: AP:1 and 1.5, DV:0.35 and 0.7

P3 coordinates: AP:0.8 and 1.2, DV:0.3 and 0.6

For targeting of RSCtx at P2, mice were anesthetized using hypothermia-induced anesthesia for 15 min and kept on an ice-cold surface during the procedure. The pups were immobilized manually, and coordinates were visually determined, except for DV: 0.4 and 0.2 from the surface of the skull.

### ICV injections

Postnatal day 0 (P0) mice were manually injected in the ventricles with 1μl of AAV9-hDlx-Flex-tdTomato (Addgene 83894) in each hemisphere after hypothermia-induced anesthesia for 8 min.

### Electrophysiology recordings in acute slices

All recordings were performed in young adults wild-type C57BL/6J mice at the VIB-KU Leuven Center for Neuroscience Technologies, Electrophysiology Unit. Briefly, mice were anesthetized with isoflurane and decapitated to prepare acute 300-μm-thick parasagittal brain slices on a Leica VT1200 vibratome. Slicing was performed in sucrose-based cutting solution (artificial cerebrospinal fluid, aCSF): 87 mM NaCl, 2.5 mM KCl, 1.25 mM NaH2PO4, 10 mM glucose, 25 mM NaHCO3, 0.5 mM CaCl2, 7 mM MgCl2, 75 mM sucrose, 1 mM kynurenic acid, 5 mM ascorbic acid and 3 mM pyruvic acid (pH 7.4 with 5% CO2/95% O2). Slices were allowed to recover at 34 °C for 30 min, and then maintained at room temperature in the same solution for at least 30 min before using. Pipettes were pulled on a horizontal micropipette puller (Sutter P-1000) and resistances ranged from 3 to 4 MΩ.

For intrinsic properties, sections were perfused at 1–2 ml/min with aCSF consisting of: 124 mM NaCl, 2.5 mM KCl, 1.2 mM NaH2PO4, 24 mM NaHCO3, 5 mM HEPES, 12.5 mM glucose, 2 mM MgSO4.7H2O, 2 mM CaCl2.2H2O and 0.05 mM picrotoxin bubbled continuously with 95% O2 and 5% CO2 and used a potassium gluconate-based internal solution: 135 mM potassium gluconate, 4 mM KCl, 2 mM NaCl, 10 mM HEPES, 4 mM EGTA, 4 mM Mg ATP and 0.3 mM Na GTP, adjusted to pH 7.25 and 295 mOsm.

Somatic current injections of 20 pA steps starting at −50 until 530 pA, from V_rest_ were used for action potential profiling. Whole-cell current clamp recordings were done at 34 °C and obtained from cell bodies that were within +/- 100 μm of the SR/SLM border visualized under the microscope. Data was collected with pCLAMP 10. Currents were sampled at 20 kHz and stored after 3 kHz low-pass Bessel filtering (Molecular Devices DigiData 1440A and Multiclamp 700B). Before analysis, the data were low-pass filtered at 1 kHz.

### Patch-seq

Before recordings we cleaned all surfaces, including recording electrodes, with RNase Away solution (Molecular BioProducts). To increase the yield of mRNA from the recorded cells, we used a small volume of recording intracellular solution (approximately 1 μl). After recordings, the cytosol of the cell was slowly aspirated via the glass pipette used for recording, with gentle negative pressure. Upon complete aspiration of the cytosol, the pipette was quickly removed from the tissue by retracting the manipulator, and the content collected to a 200 μl RNAse free PCR tube, by gently breaking the tip of the pipette at the bottom of the tube. The tube contained the lysis buffer mix Triton X-100 at 0.2% and RNase inhibitor at 1U/ μl. Collected samples were stored at -80°C until being processed.

#### cDNA library preparation

To begin with sample processing, tubes were removed from the -80°C and kept in dry ice. 2.1 µl of the oligo mix: dNTPs (10 mM each) at final concentration of 5 mM and 2.5 µM oligo-dT (/5Biosg/AAGCAGTGGTATCAACGCAGAGTACT30VN-3′) was added to each tube, and immediately placed in the Thermocycler for 3 minutes at 72°C. Next, reverse transcriptase and terminal transferase were performed using Locked nucleic acid (LNA) containing template switch oligo (5Biosg/AAGCAGTGGTATCAACGCAGAGTACATrGrG+G) (Qiagen). 5.7µl of a mix containing: 1x Superscript II buffer, 1M Betaine, 10 U/µl Superscript II RT, 5 mM DTT, Nuclease-free sterile water, 1U/µl RNase inhibitor, 1µM LNA-TSO and 6mM MgCl2, was added to each tube and placed on a thermocycler using the following conditions: 42°C for 90 min, followed by 10 cycles of 50°C for 2 min and 42°C for 2 min, with a final extension step at 70°C for 15 min.

After that, 15µl of Template switch mix containing: 1x KAPA HiFi HotStart Mix, 0.1µM IS Primers (/5Biosg/AAGCAGTGGTATCAACGCAGAGT) and Nuclease-free sterile water was added to each tube placed on a thermocycler using the following conditions: 98°C for 3 min, followed by 17 cycles of 98°C for 20 sec, 67°C for 15 sec, and 72°C for 6 min, with a final extension step at 72°C for 5 min. The PCR product was purified using AMPure XP Beads on a 0.7:1 ratio and quality control checked at the Bioanalyzer.

#### Sequencing library preparation

The purified cDNA was subsequently processed for sequencing library preparation using a modified Nextera XT tagmentation protocol. cDNA concentration was first normalized to 50 pg/µl, ensuring that no more than 1 ng total cDNA was used per reaction. For tagmentation, 1.25 µl of normalized cDNA was mixed with 3.75 µl of tagmentation mix containing 1× Tagment DNA Buffer (Illumina) and Amplicon Tagment Mix, resulting in a final reaction volume of 5 µl. Samples were incubated at 55°C for 6 min and then held at 4°C. Following tagmentation, reactions were briefly centrifuged and incubated at room temperature after addition of 1.25 µl Neutralization Buffer (NT buffer, Illumina) for 5 min.

Indexed sequencing libraries were generated by PCR amplification using Nextera index primers. Briefly, 1.25 µl each of N7xx and N5xx index primers were added to the neutralized tagmentation reaction together with 3.75 µl Nextera PCR Master Mix, yielding a final reaction volume of 12.5 µl. Libraries were amplified using the following thermocycler conditions: 72°C for 3 min, 95°C for 30 sec, followed by 13 cycles of 95°C for 10 sec, 55°C for 30 sec, and 72°C for 30 sec, with a final extension step at 72°C for 5 min. PCR products were purified using AMPure XP magnetic beads (Beckman Coulter) at a 1:1 bead-to-sample ratio.

Library quality and fragment size distribution were assessed using a Bioanalyzer (Agilent Technologies), and DNA concentration was measured using a Qubit fluorometer (Thermo Fisher Scientific). Final libraries were stored at -20°C until sequencing.

Sequencing libraries were submitted to the VIB Nucleomics Core facility for next-generation sequencing. A total of 200 individually indexed libraries were quantified using a Qubit fluorometer (Thermo Fisher Scientific), and library fragment size distribution was assessed for a subset of 21 libraries using a Bioanalyzer (Agilent Technologies). Libraries were pooled at equimolar concentrations assuming an average fragment length of 390 bp. Sequencing was performed on an Illumina NextSeq500 platform using a High Output v2.5 75-cycle kit to generate single-end 75 bp reads, yielding approximately 400 million reads in total. Raw sequencing data were demultiplexed and delivered through the Illumina BaseSpace platform.

### Data processing and analysis

#### Pre-processing

Reads from the four NextSeq500 lanes were merged per cell. Read quality was assessed with FastQC and MultiQC. Reads were aligned to the mouse reference genome mm10 (primary assembly) using STAR v2.7. Reads were counted per gene with featureCounts (Subread v2). Gene IDs were converted to gene symbols, and counts for duplicated symbols were summed. Raw gene expression data was processed and analyzed using the Seurat package (v5) in R [47]. A Seurat object was constructed with a minimum feature threshold of 500 genes per cell. Cells with more than 20% mitochondrial reads were removed.

#### Transcriptomic alignment of Patch-seq INs to a hippocampal reference atlas

Patch-seq cells were aligned to t-SNE coordinates of the Harris et al. scRNA-seq atlas of hippocampal inhibitory neurons [20,47]. Raw count matrices were loaded from the published expression table, together with pre-computed t-SNE coordinates and cluster annotations. Raw Patch-seq counts were used as the query input. Informative genes were selected from the Harris reference using a dispersion-based criterion: genes were retained if their zero-rate (fraction of cells with counts ≤ 3) exceeded an exponential function of their mean log2 expression, thereby selecting genes that are highly but not ubiquitously expressed. Patch-seq cells were projected onto the Harris t-SNE embedding using a correlation-based nearest-neighbour approach. First, the intersection of genes between the Harris reference (restricted to the selected informative genes) and the Patch-seq dataset was identified. Both count matrices were log2(count + 1) transformed. For each Patch-seq cell, Pearson correlation coefficients were computed against all Harris reference cells across the common gene set, and the 10 most correlated reference cells (k = 10 nearest neighbors) were identified. The Patch-seq cell’s t-SNE coordinates were then assigned as the median of the t-SNE positions of those 10 neighbors, yielding a single projected position per cell.

Patch-seq cells were annotated to Harris inhibitory neuron subtypes directly in gene expression space using map_to_clusters(). For each Harris cluster, a mean expression profile (centroid) was computed by averaging log2(count + 1) values across all reference cells belonging to that cluster. Each Patch-seq cell was then assigned to the cluster whose centroid showed the highest Pearson correlation with the Patch-seq cell’s expression profile, using the highly informative genes. Following annotation, Patch-seq cells that did not fall close to a Harris cluster on visual inspection of the t-SNE were marked as “undefined”.

#### Visualisations

Harris t-SNE with Patch-seq embeddings: Using R’s ggplot(), the full Harris reference embedding was rendered as a scatter plot color-coded by cluster. Patch-seq cells were overlaid as distinct markers at their projected t-SNE coordinates. Trackplots and dotplots of gene expression were generated in Scanpy4: raw counts were library-size normalized to 10,000 counts per cell and log-transformed using sc.pp.normalize_total() and sc.pp.log1p().sc.pl.tracksplot().

#### Electrophysiology data

Electrophysiological recordings were analyzed with Python using the electrophysiology feature extraction library (eFEL) [49] and custom routines. In particular, the eFEL was used to extract basic features such as resting membrane potential and membrane time constant and to detect the occurrence times of APs and their features (i.e., amplitude, half-width, threshold, and AHP amplitude, see Figs 1E and S1). The input resistance of the cells was computed by fitting a first-degree polynomial to the values of voltage deflection obtained by injecting various steps of subthreshold DC current, typically in the range [-100,100] pA in 10 – 20 pA steps. Current-frequency curves were computed by fitting the firing rate in response to the injection of suprathreshold steps of DC current with a power-law function of the form *f*(*I*) = *aI^b^* + *c*, where *a*, *b* and *c* are parameters fitted with the SciPy [50] function curve_fit. The rheobase current *I*_0_ was defined as the minimum current that produced repetitive firing and the slope of the f-I curve was computed as *ba*(2*I*_0_)*^b^*^−1^, i.e., at twice the rheobase value. Finally, the accommodation index *A* was computed by the eFEL as

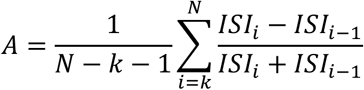

where ISI is the inter-spike interval, *N* is the total number of ISIs in a recording and *k* = 2 is the number of ISIs that were discarded in the computation. Electrophysiology readouts between annotated cell types were compared using Mann–Whitney tests, using the Holm method for multiple-testing correction.

### MERFISH spatial transcriptomics

#### Gene panel design

A gene panel of 400 genes was designed to include broad cell-type markers, inhibitory markers (based on Fig. 3 from Harris et al.) [15], differentially expressed genes from the Cck.Cxcl14 and Ntng1.Rgs10 clusters (based on S3Fig and S1.Text from Harris et al.) and genes encoding CSPs that displayed some expression in the Harris dataset. CSP-encoding genes were scanned based on current literature and using the amiGO 2 database. Probes were designed by Vizgen [51]. Due to technical limitations, we had to remove several highly expressed genes, leading to the final 368 gene panel: 18 cell type markers, 24 inhibitory markers, 70 differentially expressed genes from the Cck.Cxcl14 and Ntng1.Rgs10 clusters and 256 CSP genes.

#### Tissue processing

Three littermates from Cxcl14.Cre strain were euthanized at P10, and brains were dissected and frozen in optimal cutting temperature (O.C.T.) compound (4583, Sakura Finetek). Cryosections (10 μm) were sliced on a CM3050 S cryostat (Leica). Three non-consecutive cryosections were mounted on round functionalized coverslips (Vizgen). Tissue below the hippocampus was carefully cut off before mounting, allowing more cryosections to fit in the imageable area. A hexagonal plastic gasket was used to mark the imageable area. After 5 minutes at -20°C, coverslips were placed into a 60mm petri dish, with the tissue slice facing up. Coverslips were incubated in 4% PFA for 15 minutes at room temperature and washed twice with PBS. Then, coverslips were incubated in 70% ethanol for 5 minutes at room temperature. Lastly, coverslips were transferred each into a polytube bag (Vizgen) filled with 70% ethanol and sealed for transportation to Vizgen. During the process, areas and tools were decontaminated using RNAseZap (ThermoFisher AM9782). All solutions used were RNase-free.

### Data analysis

#### Preprocessing

Raw data was provided by Vizgen including cell segmentation MERSCOPE analysis pipeline. Data were exported from MERSCOPE Visualizer as HDF5/AnnData-derived files after manual selection of region of interest (from CA1 SP to DG granule layer, excluding CA3). Seurat objects (v5) were generated from these files, one per animal, excluding cells outside the selected hippocampal regions [47]. The three datasets were then merged, yielding 52,669 cells with 462 features (368 features plus blank probes). For QC, total RNA counts, detected features, blank fraction, and cell volume were visualized across samples. Low-quality cells were removed using thresholds of nCount_RNA >= 30, nFeature_RNA >= 20 and blank_frac < 0.2. Additional filtering was performed to exclude potential merged cells with both high volume and high RNA counts, as well as small low-count debris. Data were normalized using Seurat’s NormalizeData, and all non-blank genes were used as variable features. Data were scaled while regressing out cell volume and total RNA counts, followed by PCA, nearest-neighbour graph construction, clustering and UMAP visualization. Low-quality clusters were removed through iterative rounds of reclustering and marker-gene inspection, ending with 42100 cells.

#### Annotation

Cell classes were assigned based on the expression of canonical marker genes.

#### Reclustering of inhibitory neurons

Inhibitory neurons were then subsetted from the refined dataset and re-analysed separately using Seurat. Raw-count-based excitatory and astrocytic contamination scores were calculated from Slc17a7 and astrocyte marker genes (Gfap, Aqp4, Aldh1l1), log-transformed, and regressed out together with total RNA counts and cell volume during scaling. Marker genes for inhibitory subclusters were identified using FindAllMarkers(logfc.threshold = 0.5, only.pos = TRUE, min.pct = 0.3), and one low-quality cluster was removed before final visualization. To further assess whether CSP genes could resolve inhibitory neuron heterogeneity, inhibitory cells were also reclustered using a restricted feature set of genes encoding proteins secreted, membrane-attached or transmembrane, excluding glial markers. This resulted in a reduced feature set of 263 genes, which was used as the variable features for a second inhibitory-neuron analysis.

### scRNA sequencing

#### Tissue preparation

The neuronal extraction protocol was adapted from published work from our institute [52,53]. Each brain was immediately immersed in ice-cold carbogenated artificial cerebrospinal fluid after decapitation (Table S3). The brains were sliced in a precooled brain matrix (AgnThos, 69-2165-1) to 500-μm sections, and the hippocampi were dissected and cut into pieces in a petri dish containing ice-cold artificial cerebrospinal fluid supplemented with actinomycin D (Merck, A9415-2MG) at 1μg/ml while carbogenating. For P3, tissue from 4 pups was pooled together (3 males, 1 female). For P10 and P5, tissue from 4 animals was pooled together, 2 males, 2 females. Immediately after dissection, the tissue pieces were transferred into an enzymatic solution containing Pronase Protease (VWR 537088, 2 mg/ml), diluted in digestion medium (Table S3), and supplemented with 2% trehalose (Sigma-Aldrich, T0167) and actinomycin D. Tissue from the first dissected brains was kept in this solution on ice with carbogenation. When the tissue collection was completed, the enzymatic solution was placed in a 34°C water bath while carbogenating and incubated for 10 min, followed by the addition of 125 µl of 10mg/ml DNAse (Thermo Fisher Scientific, 10649890) and 250µl of 10mg/ml liberase (Sigma). After digestion, the enzymatic solution was removed, and tissue pieces were washed with 10 ml of digestion medium (ice-cold and carbogenated). The tissue was mechanically triturated using a volumetric pipette attached to a 100-1000µl tip in 1 ml of Opti-MEM/trehalose solution (Table S3) layered on top of a 20% Percoll solution (Table S3), and spun at 700g for 10 min at 4°C. When all solution and debris were removed, the cell pellet was resuspended in 500 μl of Opti-MEM/trehalose solution (table S3).

#### Single-cell isolation and library preparation

tdTomato^+^ and DAPI-negative cells were isolated by FACS (fluorescence-activated cell sorting) with a flow cytometer (BD Aria Fusion) using a 100 µm nozzle, about 3000-5000 events per second, and collected directly in 4μl of 10X RT Reagent E in an individual well of a 96-well plate.

The cell number and viability were assessed using the acridine orange/propidium iodide stain on the LUNA cell counter (Logos Biosystems). Only samples with cell viability of at least 80% were processed for sequencing. The samples, together with the remaining reverse transcription reaction mixture, were loaded onto a custom in-house HyDrop microfluidic droplet generator [54]. Briefly, the single-cell barcoding mix containing the cell suspension and barcoding hydrogel beads were loaded separately into mineral oil-backed syringe tips. Droplet generation oil was loaded into a third syringe. The syringes were mounted onto independent syringe pumps on the ONYX Droplet Platform (Atrandi Biosciences) and connected to a custom-fabricated HyDrop microfluidic chip [54]. Monodisperse droplets containing single cells and barcoding beads were generated by flow focusing within the microfluidic device.

Single cell RNA-seq libraries were prepared using manufacturer’s recommendations (Single cell 3’ reagent kits v3.1 user guide; CG000204 Rev D), and at the different check points the library quality was assessed using Qubit (ThermoFisher) and Bioanalyzer (Agilent). For a targeted sequencing saturation of 50-60%, sequencing was performed at a depth of 30,000 - 60,000 reads per cell and single cell libraries were sequenced on MGISEQ-2000 sequencing platform using a paired-end sequencing workflow and the recommended 10X; v3.1 read parameters (28-8-0-91 cycles).

10X libraries that were sequenced on MGISEQ-2000 sequencing platform were sequenced at the MGI Hong Kong SAR sequencing facility. Before being able to sequence on the MGI platforms, the 10X libraries needed to undergo a conversion step using MGIEasy Universal DNA Library Prep kit. Briefly, the final 10X sequencing libraries were circularized using the splint-ligation step and the circularized libraries were converted to single stranded DNA copies. DNA nanoballs were prepared from the circularized ssDNA using Rolling Circle Amplification (RCA). DNB libraries generated were then flown through the patterned flow cell of the MGISEQ-2000RS High-Throughput Sequencing kit. A custom sequencing recipe of paired end read of 28 bps (read 1), 100 bps (read 2), 10 bps (index 1) and 10 bps (index 2) was used for sequencing

#### Data analysis

##### Preprocessing

scRNA-seq data was mapped to the mm10 genome using cellranger-9.0.1. Quality control and downstream analysis were performed using Seurat in R[47]. Cells from each sample (P3, P5 and P10) were initially filtered to retain cells with at least 500 detected features. Cells with >20% mitochondrial transcripts were excluded and cells with >5% were additionally flagged in metadata for QC tracking. Ambient RNA contamination was corrected separately for each sample using SoupX [55], using sample-specific raw and filtered 10x matrices and cluster assignments from an initial merged Seurat analysis to estimate contamination profiles. SoupX-corrected count matrices were then converted back into Seurat objects, re-filtered for mitochondrial content, and doublets were identified independently for each sample using scDblFinder following standard Seurat preprocessing [56]. Cells classified as singlets were retained and merged across samples, resulting in 14,182 cells. The merged object was normalized, variable features were identified, and data were scaled while regressing out mitochondrial percentage and number of detected features. Dimensionality reduction was performed by PCA, followed by batch correction across samples using Harmony. Clustering was performed on the Harmony embedding using shared nearest-neighbour graph construction and Louvain clustering, with UMAP used for visualization. Clusters corresponding to non-inhibitory neurons and low-quality clusters were removed through iterative rounds of reclustering, marker-gene inspection and Harmony correction[27], ultimately retaining a refined population of 8,187 cells.

##### Differential gene expression analysis

DEGs among clusters were identified using Seurat’s FindAllMarkers (positive markers only) function with logfc.threshold = 1 and min.diff.pct = 0.30, and only.pos = TRUE settings. Gene ontology enrichment analysis of DEGs was performed using clusterProfiler with mouse gene annotations from org.Mm.eg.db. DEGs were considered CSP genes if the proteins encoded were transmembrane, membrane-attached, or secreted, based on UniProt entry.

##### Mapping to existing datasets

Our dataset was mapped to the Whole Mouse Brain reference (CCN20230722) [32] using MapMyCells selecting the hierarchical mapping algorithm. All cell identity annotations from the adult reference atlas were considered, without restricting mapping by brain region or cell class. Class-, subclass-, supertype-, and cluster-level annotations, together with their associated bootstrap confidence scores, were added to the Seurat metadata. Annotations with bootstrap probabilities ≥ 0.8 were retained as high-confidence assignments, whereas lower-confidence assignments were excluded from river plot visualizations. In addition, Seurat label transfer was performed to map cells to the Harris et al. dataset [15] using the FindTransferAnchors() and TransferData() functions [47]. Predicted cluster identities and maximum prediction scores were added to the query metadata, and predictions with scores <0.5 were classified as unassigned and excluded from river plot visualization. Streams with less than 20 cell were excluded from any river plot visualization.

### Single-molecule fluorescent in situ hybridization

#### Images in Fig. 2

smFISH was performed in 10 μm brain slices from WT mice mounted on SuperFrost Ultra Plus adhesion slides (Thermo Fisher Scientific). Slices were stained using the RNAscope Multiplex Fluorescent Detection Kit v2 (ACDbio, 323110) following the manufacturer’s protocols. Briefly, sections were treated for 8-10 min with a hydrogen peroxide solution and 12 min with protease IV, with 1× phosphate-buffered saline (PBS) washing in between. RNAscope probes were hybridized at 40°C for 2 hours, and subsequently, a horseradish peroxidase signal was developed using OPAL dyes according to the manufacturer’s instructions. Cxcl14 (459741), Cck (402271) and Gad2 (439371) probes were used combined with Opal 520 dye (1:1000).

#### Images in Fig. 4

smFISH was performed in 60 μm brain slices from Cxcl14.Cre mice mounted on SuperFrost Ultra Plus adhesion slides (Thermo Fisher Scientific), and dried at 40°C around 10 mins depending on age. Slices were stained using the RNAscope Multiplex Fluorescent Detection Kit v2 (ACDbio, 323110) following the manufacturer’s protocols, as described previously. *Ntng1* (488871-C3), *Rgma* (464141-C2), *Ibsp* (415501-C2), *Tmem132c (*591841-C2) and *Scube1 (*488131) probes were used combined with Opal 520 dye (1:1000).

### Tissue processing and immunohistochemistry (IHC)

Mice were transcardially perfused with 10 ml of 4% PFA under non-reversible anaesthesia. Brains were dissected and post-fixed overnight in PFA, washed in PBS and sliced in 60 μm-thick sections by vibratome. Sections were blocked in 10% normal horse serum, 0.5 M glycine, and 0.5% Triton X-100 in PBS before incubation with antibodies in 5% normal horse serum and 0.5% Triton X-100 in PBS with extensive washing between primary and secondary antibodies (Table S4). Sections were counterstained with DAPI before mounting on microscope slides with Mowiol.

### Retrobead-positive cell quantification across postnatal ages

Slices from Cxcl14.Cre brains were imaged using a Zeiss Cell Discoverer 7 and images were analysed using ImageJ. Quantification was performed manually; five or more retrobeads dots close together were considered a retrobead-positive neuron. Areas were calculated by selecting SR and SLM (polygon selection function) and using Analyze > Measure function. Values per animal correspond to the mean across 2-4 slices. Sample size per group ranged from 3 to 6 animals per timepoint. Graphs and statistical analysis were performed using R

### Quantification of *Ntng1* expression and SYT6 presence in RSCtx-projecting border INs

Slices from Cxcl14.Cre brains were imaged using a Zeiss Cell Discoverer 7, and images were analysed using ImageJ. Quantification was performed manually; 5 or more retrobead dots close together were considered a retrobead-positive neuron, while cells containing 5 or more RNAscope dots were considered positive for *Ntng1*. Sample size was 4 animals, and the values per animal correspond to the mean between 2-4 slices.

### Quantification of CSP enrichment in RSCtx-projecting vs SYT6+ border INs

Images were taken with a Nikon Spinning Disk and analysed using ImageJ. Quantification was performed manually and included the number of neurons positive for retrobeads, retrobead + CSP gene, SYT6, and SYT6 +CSP gene. NeuN+ cells containing 3 or more retrobead dots were considered retrobead-positive, while cells containing 5 or more RNAscope dots were considered positive for the respective transcript. Afterwards, the proportions Gene+ Retrobeads+/Retrobeads+ and Gene+ SYT6+/SYT6 were calculated. Graphs were plotted using R.

## ACKNOWLEDGEMENTS

We thank De Wit lab members for helpful discussion and comments. We are grateful to F. Ferrero Restelli for help with single-cell and spatial transcriptomics analysis. We thank VIB Flow Core, VIB/CBD Bioimaging Core, VIB Nucleomics Core, and VIB Single Cell Core for access to instruments and technical support.

## Data availability

The data generated in this study are available in the GEO database. The scRNAseq dataset is available under accession number GSE349671, the Patch-seq dataset is available under accession number GSE350601, and the spatial transcriptomic dataset is available under accession number GSE349320.

## SUPPLEMENTARY FIGURE LEGENDS

**Supplementary Fig. 1. Electrophysiological characterization of Patch-seq CA1 border INs.** A-E) Whole-cell current clamp recordings of intrinsic properties of CA1 border INs in acute hippocampal slices from young adult wild-type mice. Box plots show additional electrophysiological properties of Patch-seq cells assigned to the Ntng1.Rgs10 and Cck.Cxcl14 groups. Individual data points shown with bars represent mean ± SEM (*p < 0.05); Ntng1.Rgs10: n=32 cells; Cck.Cxcl14: n=37 cells. Abbreviations: AP, action potential; AHP, afterhyperpolarization.

**Supplementary Fig. 2. Spatial distribution and CSP-based clustering of CA1 border INs populations.** (A) UMAP visualization of all cells detected in the spatial transcriptomics dataset from ROI, colored by cluster.

(B) Dot plot showing expression of canonical marker genes used for cell-class annotation. Dot size indicates the percentage of cells expressing each gene, color indicates average expression.

(C) Spatial distribution of IN clusters in caudal hippocampal sections. Right side of image corresponds to proximal CA1.

(D) Dot plot showing expression of selected Cck.Cxcl14- and Ntng1.Rgs10-associated markers from the Harris dataset in spatial transcriptomics clusters identified as CA1 border INs.

(E) UMAP visualization of INs after clustering using only CSP genes, colored by the original cluster identities.

(F) Dot plot showing expression of selected Cck.Cxcl14- and Ntng1.Rgs10-associated markers in CA1 border IN clusters obtained after clustering using only CSP genes.

**Supplementary Fig. 3. Strategy to generate the *Cxcl14*-Cre line.**

**Supplementary Fig. 4 Quality control and marker-gene expression for postnatal scRNA-seq clusters.** (A) UMAP visualization of cells from the postnatal scRNA-seq dataset colored by sample. (B) Bar plot showing the sample composition of each cluster. (C) UMAP visualization showing the number of detected genes per cell. (D) UMAP visualization showing the percentage of mitochondrial transcripts per cell. (E) Feature plots showing expression of selected genes used to annotate postnatal *Cxcl14*-lineage hippocampal IN clusters. Genes shown correspond to the first and second terms used in cluster names.

**Supplementary Fig. 5 Mapping of the postnatal *Cxcl14*-lineage hippocampal IN dataset to adult reference datasets.** (A) UMAP visualization of MapMyCells class assignments. (B) River plot showing the relationship between scRNA-seq clusters and Allen Brain Cell Atlas subclasses. Cells with bootstrap probability below 0.8 were not used. Flows containing fewer than 20 cells are not shown. (C) UMAP visualization showing labels transferred from the Harris dataset. (D) River plot showing the relationship between scRNA-seq clusters and Harris dataset labels. Cells with a probability below 0.5 were not used. Flows containing fewer than 20 cells are not shown.

**Supplementary Fig. 6. Temporal dynamics of CSP DEGs in CA1 border INs.**(A) Dot plot showing top CSP genes whose expression tends to decrease over time. Dot size indicates the percentage of cells expressing each gene, colour indicates average expression. (B) Dot plot showing Cdh13 whose expression tends to increase over time. Dot size indicates the percentage of cells expressing each gene, and colour indicates average expression. (C) Dot plot showing top CSP genes whose expression presents opposing trends over time depending on the CA1 border IN cluster. Dot size indicates the percentage of cells expressing each gene, colour indicates average expression.

**Supplementary Fig. 7. Retrobead labeling in CA1 following RSCtx L1 injection at different postnatal ages.** (A) Representative images showing retrobead labeling (green) two days after retrobead injection in RSCtx L1 at the indicated postnatal ages. Dotted circles highlight retrobead labeling in the thalamus. Dotted boxes represent from which the zoomed SR/SLM border crop was obtained. Arrowheads point to retrobead-labelled cells.

**Supplementary Fig. 8. Spatial distribution of CSP gene expression in the postnatal hippocampus.**(A) Representative hippocampal images showing CSP gene expression (magenta) and NeuN immunohistochemistry (white).

## SUPPLEMENTARY TABLES

**Supplementary Table 1. Spatial transcriptomics gene probe list**

**Supplementary Table 2. Detailed mapping of the postnatal *Cxcl14*-lineage hippocampal IN dataset to adult reference datasets.** This workbook contains the cell-type mapping results underlying the river plot shown in Fig. S5. The “Summary cluster ID” worksheet summarizes the top-matching annotation and the percentage of low-confidence assignments for each cluster. Cluster colours are consistent with those used for the corresponding streams in the river plots in Fig. S5. Additional worksheets provide the complete distribution of cell assignments at different annotation levels: Harris broader class, Harris cluster, MapMyCells subclass, MapMyCells supertype, and MapMyCells cluster. For each annotation level, the frequency and percentage of cells assigned to each reference cell type are reported, together with the percentage calculated among successfully mapped cells.

**Supplementary Table 3. Composition of solutions used for single-cell preparation.** Composition, final concentrations, and supplier identifiers of solutions used for tissue processing and single-cell preparation.

**Supplementary Table 4. Primary antibodies used in this study.** Primary antibodies used for IHC, including working dilution, manufacturer, and product identifier.

## REFERENCES

1. Christensen R, Shao Z, Colón-Ramos DA. The cell biology of synaptic specificity during development. Curr Opin Neurobiol. 2013;23: 10.1016/j.conb.2013.07.004. doi:10.1016/j.conb.2013.07.004

2. Sanes JR, Zipursky SL. Synaptic Specificity, Recognition Molecules, and Assembly of Neural Circuits. Cell. 2020;181: 536–556. doi:10.1016/j.cell.2020.04.008

3. Yogev S, Shen K. Cellular and Molecular Mechanisms of Synaptic Specificity. Annu Rev Cell Dev Biol. 2014;30: 417–437. doi:10.1146/annurev-cellbio-100913-012953

4. van Strien NM, Cappaert NLM, Witter MP. The anatomy of memory: an interactive overview of the parahippocampal–hippocampal network. Nat Rev Neurosci. 2009;10: 272–282. doi:10.1038/nrn2614

5. Pelkey KA, Chittajallu R, Craig MT, Tricoire L, Wester JC, McBain CJ. Hippocampal gabaergic inhibitory interneurons. Physiol Rev. 2017;97: 1619–1747. doi:10.1152/physrev.00007.2017

6. Tzilivaki A, Tukker JJ, Maier N, Poirazi P, Sammons RP, Schmitz D. Hippocampal GABAergic interneurons and memory. Neuron. 2023;111: 3154–3175. doi:10.1016/J.NEURON.2023.06.016

7. Booker SA, Vida I. Morphological diversity and connectivity of hippocampal interneurons. Cell Tissue Res. 2018;373: 619–641. doi:10.1007/s00441-018-2882-2

8. Kepecs A, Fishell G. Interneuron Cell Types: Fit to form and formed to fit. Nature. 2014;505: 318–326. doi:10.1038/nature12983

9. Tricoire L, Pelkey KA, Erkkila BE, Jeffries BW, Yuan X, McBain CJ. A blueprint for the spatiotemporal origins of mouse hippocampal interneuron diversity. J Neurosci. 2011;31: 10948–10970. doi:10.1523/JNEUROSCI.0323-11.2011

10. Klausberger T. GABAergic interneurons targeting dendrites of pyramidal cells in the CA1 area of the hippocampus. Eur J Neurosci. 2009;30: 947–957. doi:10.1111/j.1460-9568.2009.06913.x

11. Vida I, Halasy K, Szinyei C, Somogyi P, Buhl EH. Unitary IPSPs evoked by interneurons at the stratum radiatum-stratum lacunosum-moleculare border in the CA1 area of the rat hippocampus in vitro. J Physiol. 1998;506: 755–773. doi:10.1111/j.1469-7793.1998.755bv.x

12. Fuentealba P, Klausberger T, Karayannis T, Suen WY, Huck J, Tomioka R, et al. Expression of COUP-TFII Nuclear Receptor in Restricted GABAergic Neuronal Populations in the Adult Rat Hippocampus. J Neurosci. 2010;30: 1595–1609. doi:10.1523/JNEUROSCI.4199-09.2010

13. Klausberger T, Somogyi P. Neuronal Diversity and Temporal Dynamics: The Unity of Hippocampal Circuit Operations. Science. 2008;321: 53–57. doi:10.1126/science.1149381

14. Yamawaki N, Li X, Lambot L, Ren LY, Radulovic J, Shepherd GMG. Long-range inhibitory intersection of a retrosplenial thalamocortical circuit by apical tuft-targeting CA1 neurons. Nat Neurosci. 2019;22: 618–626. doi:10.1038/s41593-019-0355-x

15. Harris KD, Hochgerner H, Skene NG, Magno L, Katona L, Bengtsson Gonzales C, et al. Classes and continua of hippocampal CA1 inhibitory neurons revealed by single-cell transcriptomics. PLoS Biology. 2018. doi:10.1371/journal.pbio.2006387

16. De Wit J, Ghosh A. Specification of synaptic connectivity by cell surface interactions. Nat Rev Neurosci. 2016;17: 22–35. doi:10.1038/nrn.2015.3

17. Földy C, Darmanis S, Aoto J, Malenka RC, Quake SR, Südhof TC. Single-cell RNAseq reveals cell adhesion molecule profiles in electrophysiologically defined neurons. Proc Natl Acad Sci U S A. 2016;113: E5222–E5231. doi:10.1073/pnas.1610155113

18. Cadwell CR, Scala F, Li S, Livrizzi G, Shen S, Sandberg R, et al. Multimodal profiling of single-cell morphology, electrophysiology and gene expression using Patch-seq. Nat Protoc. 2017;12: 2531–2553. doi:10.1038/nprot.2017.120

19. Tasic B, Yao Z, Graybuck LT, Smith KA, Nguyen TN, Bertagnolli D, et al. Shared and distinct transcriptomic cell types across neocortical areas. Nature. 2018;563: 72–78. doi:10.1038/s41586-018-0654-5

20. Kobak D, Berens P. The art of using t-SNE for single-cell transcriptomics. Nat Commun. 2019;10: 5416. doi:10.1038/s41467-019-13056-x

21. Liodis P, Denaxa M, Grigoriou M, Akufo-Addo C, Yanagawa Y, Pachnis V. Lhx6 Activity Is Required for the Normal Migration and Specification of Cortical Interneuron Subtypes. J Neurosci. 2007;27: 3078–3089. doi:10.1523/JNEUROSCI.3055-06.2007

22. Lim L, Mi D, Llorca A, Marín O. Development and functional diversification of cortical interneurons. Neuron. 2018;100: 294–313. doi:10.1016/j.neuron.2018.10.009

23. Favuzzi E, Deogracias R, Marques-Smith A, Maeso P, Jezequel J, Exposito-Alonso D, et al. Neurodevelopment: Distinct molecular programs regulate synapse specificity in cortical inhibitory circuits. Science. 2019;363: 413–417. doi:10.1126/science.aau8977

24. Madisen L, Zwingman TA, Sunkin SM, Oh SW, Zariwala HA, Gu H, et al. A robust and high-throughput Cre reporting and characterization system for the whole mouse brain. Nat Neurosci. 2010;13: 133–140. doi:10.1038/nn.2467

25. Southwell DG, Paredes MF, Galvao RP, Jones DL, Froemke RC, Sebe JY, et al. Intrinsically determined cell death of developing cortical interneurons. Nature. 2012;491: 109–113. doi:10.1038/nature11523

26. Zheng GXY, Terry JM, Belgrader P, Ryvkin P, Bent ZW, Wilson R, et al. Massively parallel digital transcriptional profiling of single cells. Nat Commun. 2017;8: 14049. doi:10.1038/ncomms14049

27. Korsunsky I, Millard N, Fan J, Slowikowski K, Zhang F, Wei K, et al. Fast, sensitive and accurate integration of single-cell data with Harmony. Nat Methods. 2019;16: 1289–1296. doi:10.1038/s41592-019-0619-0

28. van Velthoven CTJ, Gao Y, Kunst M, Lee C, McMillen D, Chakka AB, et al. Transcriptomic and spatial organization of telencephalic GABAergic neurons. Nature. 2025;647: 143–156. doi:10.1038/s41586-025-09296-1

29. Katona L, Hartwich K, Tomioka R, Somogyi J, Roberts JDB, Wagner K, et al. Synaptic organisation and behaviour-dependent activity of mGluR8a-innervated GABAergic trilaminar cells projecting from the hippocampus to the subiculum. Brain Struct Funct. 2020;225: 705–734. doi:10.1007/s00429-020-02029-2

30. Ferraguti F, Klausberger T, Cobden P, Baude A, Roberts JDB, Szucs P, et al. Metabotropic Glutamate Receptor 8-Expressing Nerve Terminals Target Subsets of GABAergic Neurons in the Hippocampus. J Neurosci. 2005;25: 10520–10536. doi:10.1523/JNEUROSCI.2547-05.2005

31. Conrad S, Genth H, Hofmann F, Just I, Skutella T. Neogenin-RGMa Signaling at the Growth Cone Is Bone Morphogenetic Protein-independent and Involves RhoA, ROCK, and PKC *. J Biol Chem. 2007;282: 16423–16433. doi:10.1074/jbc.M610901200

32. Yao Z, van Velthoven CTJ, Kunst M, Zhang M, McMillen D, Lee C, et al. A high-resolution transcriptomic and spatial atlas of cell types in the whole mouse brain. Nat 2023 6247991. 2023;624: 317–332. doi:10.1038/s41586-023-06812-z

33. Nishimura-Akiyoshi S, Niimi K, Nakashiba T, Itohara S. Axonal netrin-Gs transneuronally determine lamina-specific subdendritic segments. Proc Natl Acad Sci U S A. 2007;104: 14801. doi:10.1073/PNAS.0706919104

34. Que L, Lukacsovich D, Luo W, Földy C. Transcriptional and morphological profiling of parvalbumin interneuron subpopulations in the mouse hippocampus. Nat Commun. 2021;12: 108. doi:10.1038/s41467-020-20328-4

35. Monnier PP, Sierra A, Macchi P, Deitinghoff L, Andersen JS, Mann M, et al. RGM is a repulsive guidance molecule for retinal axons. Nature. 2002;419: 392–395. doi:10.1038/nature01041

36. Tu CF, Yan YT, Wu SY, Djoko B, Tsai MT, Cheng CJ, et al. Domain and functional analysis of a novel platelet-endothelial cell surface protein, SCUBE. J Biol Chem. 2008;283: 12478–12488. doi:10.1074/jbc.M705872200

37. Lin YC, Sahoo BK, Gau SS, Yang RB. The biology of SCUBE. J Biomed Sci 2023 301. 2023;30: 33-. doi:10.1186/S12929-023-00925-3

38. Kottmann V, Drees P, Gercek E, Ritz U. Bone sialoprotein: a multifunctional regulator of bone remodelling and tumour progression. Bone Res 2026 141. 2026;14: 11-. doi:10.1038/s41413-025-00490-5

39. Sanchez-Pulido L, Ponting CP. TMEM132: an ancient architecture of cohesin and immunoglobulin domains define a new family of neural adhesion molecules. Bioinformatics. 2018;34: 721–724. doi:10.1093/BIOINFORMATICS/BTX689

40. Wang Y, Herzig G, Molano C, Liu A. Differential expression of the Tmem132 family genes in the developing mouse nervous system. Gene Expr Patterns. 2022;45: 119257. doi:10.1016/J.GEP.2022.119257

41. Wang X, Jiang W, Luo S, Yang X, Wang C, Wang B, et al. The C. elegans homolog of human panic-disorder risk gene TMEM132D orchestrates neuronal morphogenesis through the WAVE-regulatory complex. Mol Brain 2021 141. 2021;14: 54-. doi:10.1186/S13041-021-00767-W

42. Iannone AF, Akgül G, Zhang R, Wacks S, Hussein N, Macias CG, et al. The chemokine Cxcl14 regulates interneuron differentiation in layer I of the somatosensory cortex. Cell Rep. 2024;43: 114531. doi:10.1016/J.CELREP.2024.114531

43. Tanegashima K, Suzuki K, Nakayama Y, Tsuji K, Shigenaga A, Otaka A, et al. CXCL14 is a natural inhibitor of the CXCL12-CXCR4 signaling axis. FEBS Lett. 2013;587: 1731–1735. doi:10.1016/j.febslet.2013.04.046

44. Stumm RK, Zhou C, Ara T, Lazarini F, Dubois-Dalcq M, Nagasawa T, et al. CXCR4 Regulates Interneuron Migration in the Developing Neocortex. J Neurosci. 2003;23: 5123–5130. doi:10.1523/JNEUROSCI.23-12-05123.2003

45. Wang Y, Li G, Stanco A, Long JE, Crawford D, Potter GB, et al. CXCR4 and CXCR7 Have Distinct Functions in Regulating Interneuron Migration. Neuron. 2011;69: 61–76. doi:10.1016/j.neuron.2010.12.005

46. Challis RC, Ravindra Kumar S, Chan KY, Challis C, Beadle K, Jang MJ, et al. Systemic AAV vectors for widespread and targeted gene delivery in rodents. Nat Protoc. 2019;14: 379–414. doi:10.1038/s41596-018-0097-3

47. Hao Y, Stuart T, Kowalski MH, Choudhary S, Hoffman P, Hartman A, et al. Dictionary learning for integrative, multimodal and scalable single-cell analysis. Nat Biotechnol. 2024;42: 293–304. doi:10.1038/s41587-023-01767-y

48. Choudhary S, Satija R. Comparison and evaluation of statistical error models for scRNA-seq. Genome Biol. 2022;23: 27. doi:10.1186/s13059-021-02584-9

49. Mandge D, Tuncel A, Jaquier A, Kilic I, Damart T, Markram H, et al. eFEL: electrophysiology feature extraction library. Bioinformatics. 2026;42: btag328. doi:10.1093/bioinformatics/btag328

50. Virtanen P, Gommers R, Oliphant TE, Haberland M, Reddy T, Cournapeau D, et al. SciPy 1.0: fundamental algorithms for scientific computing in Python. Nat Methods. 2020;17: 261–272. doi:10.1038/s41592-019-0686-2

51. Moffitt JR, Bambah-Mukku D, Eichhorn SW, Vaughn E, Shekhar K, Perez JD, et al. Molecular, Spatial and Functional Single-Cell Profiling of the Hypothalamic Preoptic Region. Science. 2018;362: eaau5324. doi:10.1126/science.aau5324

52. Lorente-Echeverría B, Daaboul D, Vandensteen J, Marcassa G, Naert W, Vandenbempt J, et al. A dynamic gene regulatory code drives synaptic development of hippocampal granule cells. Sci Adv. 2025;11: eadx5140. doi:10.1126/sciadv.adx5140

53. Liu M, Restelli FF, Micoli E, Barbiera G, Moors R, Nouboers E, et al. Single-cell transcriptome of developing inhibitory neurons reveals expanding and contracting modes of diversification. bioRxiv; 2025. p. 2025.02.19.636192. doi:10.1101/2025.02.19.636192

54. De Rop FV, Ismail JN, Bravo González-Blas C, Hulselmans GJ, Flerin CC, Janssens J, et al. Hydrop enables droplet-based single-cell ATAC-seq and single-cell RNA-seq using dissolvable hydrogel beads. Barkai N, Mulder K, editors. eLife. 2022;11: e73971. doi:10.7554/eLife.73971

55. Young MD, Behjati S. SoupX removes ambient RNA contamination from droplet-based single-cell RNA sequencing data. GigaScience. 2020;9: giaa151. doi:10.1093/gigascience/giaa151

56. Germain P-L, Lun A, Meixide CG, Macnair W, Robinson MD. Doublet identification in single-cell sequencing data using *scDblFinder*. F1000Research; 2022. doi:10.12688/f1000research.73600.2

